# Asymmetric nuclear division of neural stem cells contributes to the formation of sibling nuclei with different identities

**DOI:** 10.1101/2020.08.29.272724

**Authors:** Chantal Roubinet, Ian J. White, Buzz Baum

## Abstract

Cellular diversity in multicellular organisms is often generated via asymmetric divisions. In the fly, for example, neural stem cells divide asymmetrically to generate a large self-renewing stem cell and a smaller sibling that differentiates. Efforts to understand how these different cell fates are generated have focused on the asymmetric segregation of cortically-localised transcription factors at division, which preferentially enter single daughter cell nuclei to change their fate. However, we find that the nuclear compartment in these cells remains intact throughout mitosis and is asymmetrically inherited, giving rise to sibling nuclei that differ profoundly in size, envelope composition and fate markers. These data reveal the importance of considering nuclear remodelling during stem cell divisions, and show how daughter cell fates depend on the coordination of the asymmetric inheritance of cortical fate markers with asymmetric nuclear division.

## Introduction

The division of a fly neural stem cell generates two sibling daughter cells that have distinct fates. While previous studies have largely attributed this to the asymmetric inheritance of cortically localized transcription factors, which bias gene expression in the two daughter cells causing each to assume a different fate (self-renewal *versus* differentiation), neuroblast divisions also generate cells with daughter cell nuclei that are very different in size. Here, to shed light on this process of asymmetric nuclear division, we set out to investigate how differences in sibling nuclei arise during mitotic progression, and to determine the relationship between nuclear division and asymmetric cell division.

Studies over many decades have revealed a strong correlation between interphase nuclear size, genome size and cell size over evolutionary timescales ^1^ and, in the context of development, nuclear size tends to scale with cell size ^2-4^. Although often ignored as a factor, the size of the nucleus will also depend on the process by which nuclei are formed at mitotic exit. How this works is not well understood but will depend on the system. In many eukaryotes, which exhibit a so-called “open mitosis”, the nuclear envelope is reformed anew at mitotic exit ^5-7^. By contrast, other eukaryotes (e.g. fungi and dinoflagellates) undergo a so-called “closed mitosis” in which they maintain an intact nuclear compartment and a diffusion barrier throughout ^7, 8^. Interestingly, *Drosophila* appear to lie somewhere in between these two extremes. Thus, the nuclear envelope remains partially intact in the syncytial fly embryo^9, 10^, but appears to break down during mitosis in *Drosophila* cells in culture ^11^.

Here, our goal was to determine how the nuclear compartment is remodelled during stem cell divisions to give rise to daughter cells with nuclei of distinct size and fates. Through this analysis, we find that daughter nuclei with a distinct size and molecular composition arise during the course of neuroblast division as a consequence of both asymmetric division and growth of the nuclear compartment. Strikingly, these asymmetries in nuclear size and composition persist even when nuclei enter the same cytoplasm following late division failure – leading to the generation of distinct nuclei in the same cytoplasm. Thus, in the context of neural stem cell divisions, the identity of daughter cells depends on the temporal and spatial coupling of asymmetric division, driven by a polarised actomyosin cortex, with the asymmetric segregation and growth of the nuclear compartment.

## Results

### The nuclear envelope of fly neuroblasts is maintained during mitosis and remodelled to generate two sibling nuclei that differ in size and composition

To characterize nuclear division of *Drosophila* neuroblasts, we performed time-lapse microscopy using various nuclear membrane markers in cells co-expressing the spindle marker Cherry::Jupiter. This includes CD8::GFP, a marker that labels both the plasma membrane and the nuclear envelope ^12^ (Figure 1A; yellow arrows; Supplemental Video 1), Klaroid::GFP and Klarischt::GFP (markers of the Inner Nuclear Membrane (INM) and Outer Nuclear Membrane (ONM) respectively), and the Endoplasmic Reticulum (ER)/Nuclear envelope (NE) lumenal reporter protein BIP::SfGFP::HDEL (Figure 1B, Supplemental Video 2 and data not shown). In all cases, these markers labelled a membrane that enveloped the spindle through metaphase and anaphase, before being remodeled to form two nuclei with different sizes at mitotic exit. These data show that the membrane delimiting the nuclear compartment is maintained during cell division, and retains elements of its identity (INM, ONM and ER lumen) as neuroblasts pass throughout mitosis. These observations are in keeping with data suggesting that some fly cells (e.g. in the syncytial embryo) ^10^ undergo a “semi-open mitosis”. However, the presence of a nuclear envelope does not mean that the nuclear/cytoplasmic compartment boundary is maintained during mitosis. In fact, a marker of the nucleoplasm, NLS::5GFP (a fusion of five GFPs with a nuclear localization sequence that is found in nuclei in interphase cells) was seen rapidly leaving nuclei as cells entered prometaphase, leading to uniform levels between the nuclear and cytoplasmic compartments until late telophase (Figure 1C; top panel, Figure S1A,C and Supplemental Video 3). Conversely, the cytoplasmic marker RpS13::GFP rapidly entered the nuclear compartment during prometaphase, confirming loss of nuclear/cytoplasmic diffusion barrier upon entry into mitosis (Figure 1C; bottom panel, Figure S1B,D and Supplemental Video 4).

**Figure 1:**
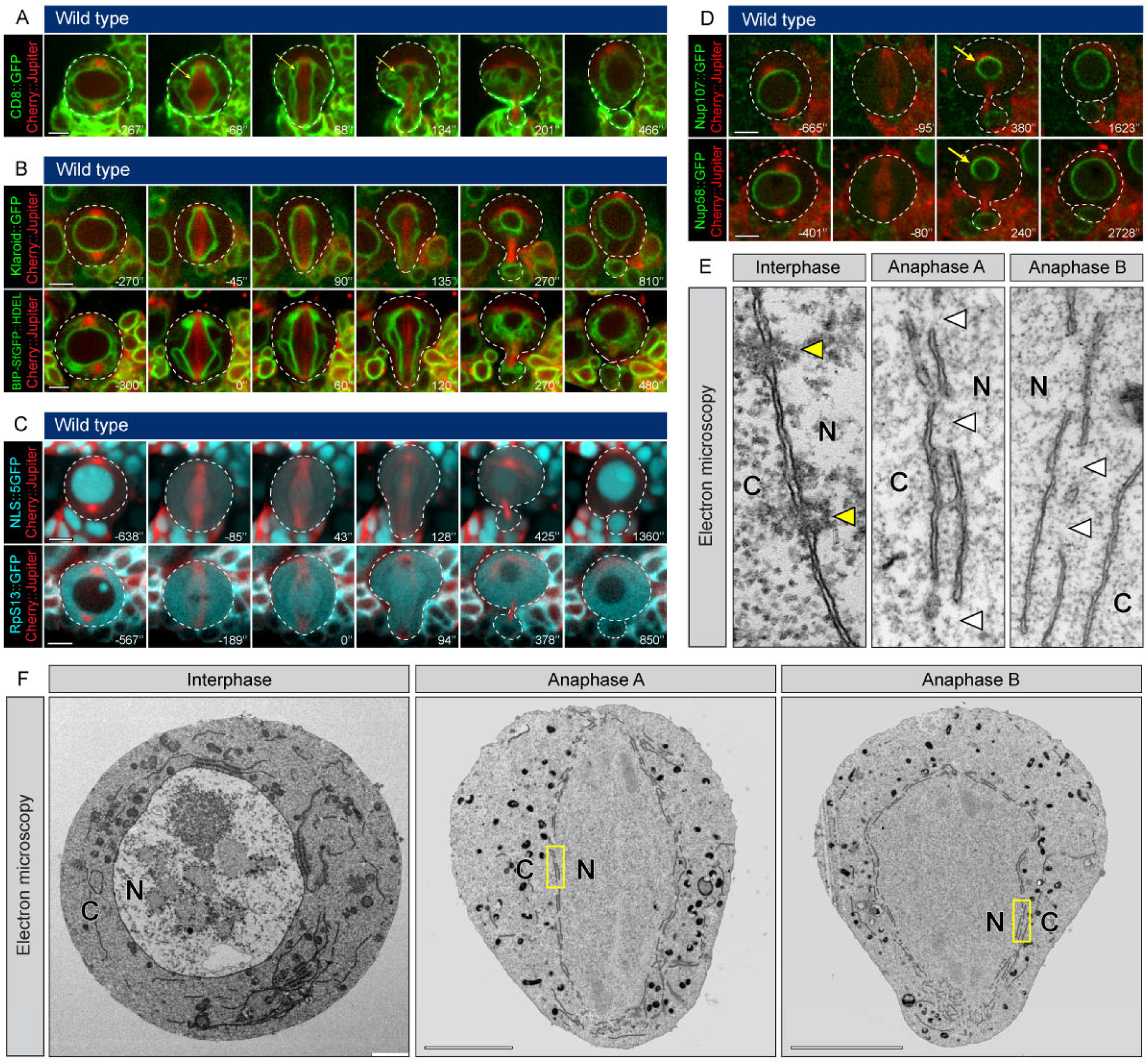
The nuclear envelope of fly neuroblasts is maintained during mitosis and remodelled to generate two sibling nuclei that differ in size and composition. (**A-D**) Mitotic neuroblasts imaged in intact brain, expressing the spindle marker Cherry::Jupiter (red) in combination with either (**A**) plasma and nuclear membranes marker CD8::GFP (green), (**B**) INM or ER marker Klaroid::GFP and BIP-SfGFP::HDEL respectively (green), (**C**) NLS::5GFP or the ribosomal subunit Rps13::GFP (blue), (**D**) the nucleoporins Nup107::GFP or Nup58::GFP (green). Yellow arrows point the higher Nucleoporin density on neuroblast nucleus. (**E, F**) Electron microscopy performed on isolated neuroblasts obtained from dissociated brains, at various cell cycle stages: interphase, anaphase A and anaphase B. (**E**) show magnified images of area marked in yellow in (**F**); yellow arrow heads show nuclear pores; white arrow heads show holes in the nuclear envelope; N = nuclear compartment; C = cytoplasm. Scale bars are 5µm.

To determine why the neuroblast nuclear compartment boundary becomes permeable to large proteins and ribosomes during mitosis, we imaged the Nuclear Pore Complexes (called NPCs), since they are responsible for the regulated trafficking of soluble proteins between the cytoplasmic and nuclear compartments. Time lapse imaging of neuroblasts expressing tagged versions of an FG-repeat containing nucleoporin (Nup58::GFP), which contributes to the maintenance of the NPCs diffusion barrier, or a nucleoporin that forms part of the structural scaffold of the outer pore (Nup107::GFP), revealed that both nucleoporins were lost from the nuclear membrane upon entry into mitosis, before being recruited back to the newly formed nuclei following mitotic exit (Figure 1D). At this stage, the recruitment of both nucleoporins to daughter nuclei was strikingly asymmetric, resulting in a higher density of NPCs within the nascent neuroblast nuclear envelope compared to that found in the nucleus of the differentiating sibling (Figure 1D; yellow arrows, Figure S1E). Importantly, these differences in envelope composition in daughter nuclei were established before the cortical release of polarity proteins (Figure S1F).

The loss of the Nup58 signal at the nuclear envelope might reflect a partial disassembly of NPCs, as has been seen in *Aspergillus nidulans* ^*13*^, or the complete loss of NPCs. Thus, to assess the nuclear membrane ultrastructure during passage through mitosis, we performed electron microscopy on isolated neuroblasts in both interphase and mitosis. This revealed that NPCs were completely disassembled during mitosis (Figure 1E; yellow arrow heads in interphase). The mitotic nuclear membrane that remained encased the spindle throughout mitosis (Figure 1F) without becoming dispersed into the cytoplasm (as occurs in most mitotic mammalian and *Drosophila* cells in culture ^11, 14^), and appeared highly fenestrated and multilayered (Figure 1E; white arrow heads & 1F). While allowing for the free movement of ribosomes (Figure 1C; bottom panel & 1E), the openings in this multilayered mitotic nuclear envelope did not appear large enough to allow the passage of larger structures. In line with this, vesicles and mitochondria remained excluded into the spindle region (Figure 1F, Figure S1G,H). These data show that *Drosophila* neuroblasts undergo a semi-closed mitosis, characterized by the complete disassembly of NPCs and the formation of a multilayered fenestrated mitotic nuclear envelope.

### A nuclear lamina is required for nuclear envelope maintenance during passage through mitosis

In a search for factors that maintain this bounding nuclear membrane in mitotic cells that have disassembled their NPCs, we focused on the Lamins. These highly conserved proteins form a physical support for the nuclear membrane in interphase, but are typically lost from mitotic nuclei in cells that undergo nuclear membrane fragmentation ^15^. When we imaged cells over-expressing GFP-tagged LamDm0, the sole B-type Lamin in the *Drosophila* genome, we saw that, while the levels of LamDm0::GFP present at the interphase nuclear envelope were visibly reduced during mitosis (1.8-fold), some LamDm0::GFP remained at the nuclear envelope throughout (Figure 2A,B,C and Supplemental Video 5). Importantly, antibody staining confirmed that this also holds for endogenous LamDm0 (Figure 2D). Furthermore, because the mitotic phosphorylation of Lamin promotes lamina disassembly in many systems ^15, 16^, we also stained tissue with an antibody that specifically recognises a form of endogenous LamDm0 that lacks phosphorylation on Serine 25. Again, this was seen forming a lamina that underlies the bounding mitotic nuclear envelope (Figure 2E).

**Figure 2:**
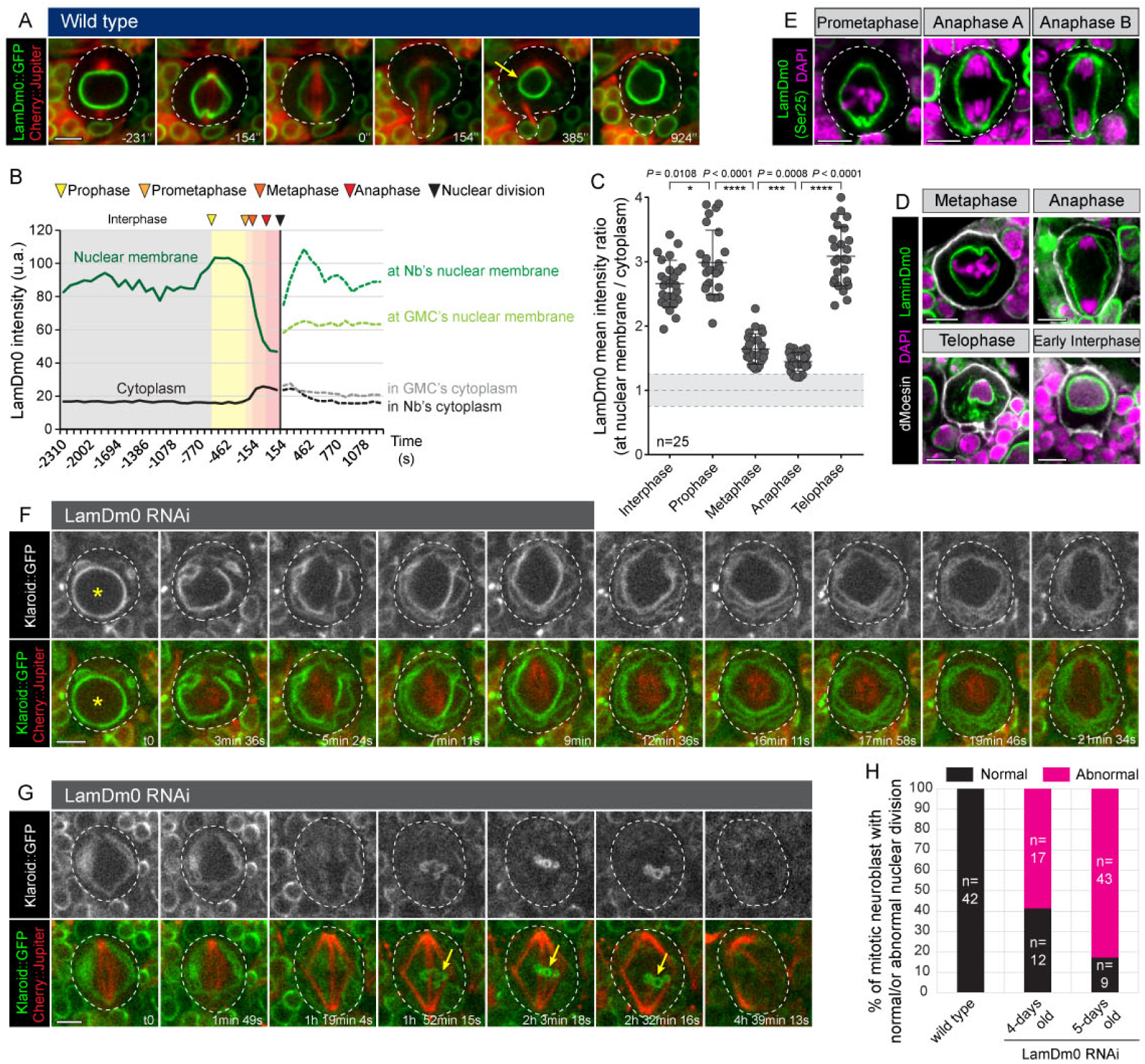
A nuclear lamina is required for nuclear envelope maintenance during passage through mitosis. (**A**) Mitotic neuroblast imaged in intact brain, expressing the lamin LamDm0::GFP (green) with the spindle marker Cherry::Jupiter (red). Yellow arrow point the higher LamDm0 density on the neuroblast nucleus. (**B**) LamDm0::GFP mean intensity is measured along the nuclear membrane (green curves) as well as in the cytoplasm (black curves) throughout the cell cycle, and plotted in the graph. The coloured areas on the graph correspond to different stages of the cell cycle. (**C**) LamDm0 mean intensity ratio between the nuclear membrane and the cytoplasm is measured at 5 different stages of the cell cycle for 25 cells. The grey area on the graph corresponds to a ratio equal to 1 ± 0.25. Bars indicate mean ± standard deviation. Asterisks denote statistical significance, derived from unpaired t tests: *: p ≤ 0.05, ***: p ≤ 0.001 and ****: p ≤ 0.0001. (**D**) Immunostaining of dividing neuroblasts for LamDm0 (green) and the cortical marker dMoesin (white). DNA is stained with DAPI (purple). (**E**) Immunostaining of dividing neuroblasts for unphosphorylated LamDm0 on Serine 25 (green). DNA is stained with DAPI (Purple). (**F**,**G**) Representative time lapses of neuroblast expressing the INM marker Klaroid (white on the top panels; green on the merges), the spindle marker Cherry::Jupiter (red on the merges) and an RNAi directed against LamDm0. Yellow stars show the presence of an intact nuclear compartment during prophase and yellow arrows show nuclear membrane recruitment around chromatin. (**H**) Quantification of mitotic neuroblasts expressing an RNAi against LamDm0 and displaying either a normal or abnormal nuclear division (e.g. dispersion of nuclear membrane fragments into the cytoplasm from metaphase onward, or rupture of the nuclear envelope). Larvae of 4 and 5 days old are dissected. Number of analysed cells = 123. In each case, data were collected from at least 3 independent experiments. Scale bars are 5µm.

To test whether the preservation of this mitotic nuclear lamina plays a role in the maintenance of a nuclear compartment by preventing nuclear envelope fragments to disperse into the cytoplasm, we used RNAi to reduce levels of LaminDm0 expression. While the nuclear envelope looks normal during interphase and prophase (Figure 2F; star), LamDm0 depletion led to dispersion of nuclear membrane fragments into the cytoplasm in 43% of cells progressing through mitosis (Figure 2F). This suggests that reducing levels of LamDm0 is sufficient to convert semi-closed mitosis of neural stems cells in one that appears open. Furthermore, the dispersion of nuclear membrane into the cytoplasm was often associated with a dramatic delay of the anaphase onset, the formation of defective spindles and a failure in chromosome segregation, culminating in the late recruitment of nuclear membrane around the metaphase plate and cell death (Figure 2G; yellow arrows). Abnormal nuclear division was seen in more than 80% of mitotic neuroblasts expressing an RNAi against LamDm0 (Figure 2H; 5-days old larvae). These data show the importance of maintaining an intact nuclear envelope during neuronal stem cell divisions, and highlight the importance of the mitotic lamina for semi-closed mitosis in fly neuroblasts. We next sought to understand how this mitotic nuclear compartment is remodelled to form two nuclei of different size.

### Asymmetric nuclear division depends on an asymmetric nuclear sealing at sites dictated by the spindle

To determine how the nuclear envelope is remodelled at mitotic exit to generate two distinct daughter nuclei, we used CD8::GFP ^12^ to study how the changes in the shape of the plasma membrane and the bounding nuclear membrane are coordinated. Both membranes developed an apico-basal asymmetry at the same time, as apical width increased relative to basal width (Figure 3A,B; compare time points −25” with 123”). Both the cell and nuclear compartment elongated at a similar rate at the onset of anaphase (Figure 3C), and both narrowed at the same position prior to cytokinesis (Figure 3D). Furthermore, cortical furrowing consistently preceded constriction of the nucleus with a short time-delay (Figure 3D; blue *versus* green arrow heads). These data suggested the possibility that the contraction of the cleavage furrow squeezes the nuclear compartment in a basally-shifted position, leading to a local rupture or fusion of nuclear membranes to generate large and small daughter nuclei. To test whether these two events (cell and nuclear division) are causally linked, as suggested by this analysis, we perturbed contraction of the cleavage furrow using the actin-poisons Latrunculin A and Cytochalasin D, or by plating cultured isolated neuroblasts on Concanavalin A-coated dishes which interferes with furrow formation by forcing cells to adhere to the dish ^17^. Although a portion of neuroblasts completed cytokinesis under these conditions, most ultimately failed to divide (Figure 3e, Figure S2A and Supplemental Video 6). Importantly, from 559 mitotic neuroblasts imaged, cytokinesis failure always yielded single cells with two nuclei rather than one (Figure S2A), even after successive cytokinesis failures (Figure S2B). Thus, nuclear division did not require the contraction of the cleavage furrow. Nor was it driven by a local rupture or fusion of nuclear membrane. Instead, live cell imaging revealed that the fenestrated mitotic nuclear envelope was re-sealed to generate the daughter nuclei, via progressive extension of the nuclear envelope (Supplemental Video 6). Interestingly, this resealing process began from the lateral membranes and was completed nearer the cell center, at sites that correspond to the extremities of the central spindle (Figure 3F; yellow arrow heads and kymograph). If, as suggested by these findings, the bundles of microtubules within the central spindle play a role in defining the sites at which sealing of the mitotic nuclear compartment occurs, it should be possible to impair nuclear division using perturbations that affect the central spindle. To test this prediction, we used different concentrations of the Aurora B-specific inhibitor Binuclein 2 ^18^ to compromise central spindle formation. In these experiments, when a central spindle was visible (Figure S2C; orange arrow), two nuclear compartments were formed, leading to the generation of binucleated cells after cleavage furrow regression (Figure S2E; black group). By contrast, loss of the central spindle (Figure S2D; orange arrow) was associated with a failure of cells to undergo both nuclear sealing and cell division, leading to the formation of cells containing a single, large nuclear compartment (Figure S2E; pink group). Consistent with these observations, RNAi-mediated silencing of Aurora B led to a similar impairment of the central spindle formation, which was associated with failed nuclear sealing and, ultimately, a failure in nuclear division (Figure 3G,H).

**Figure 3:**
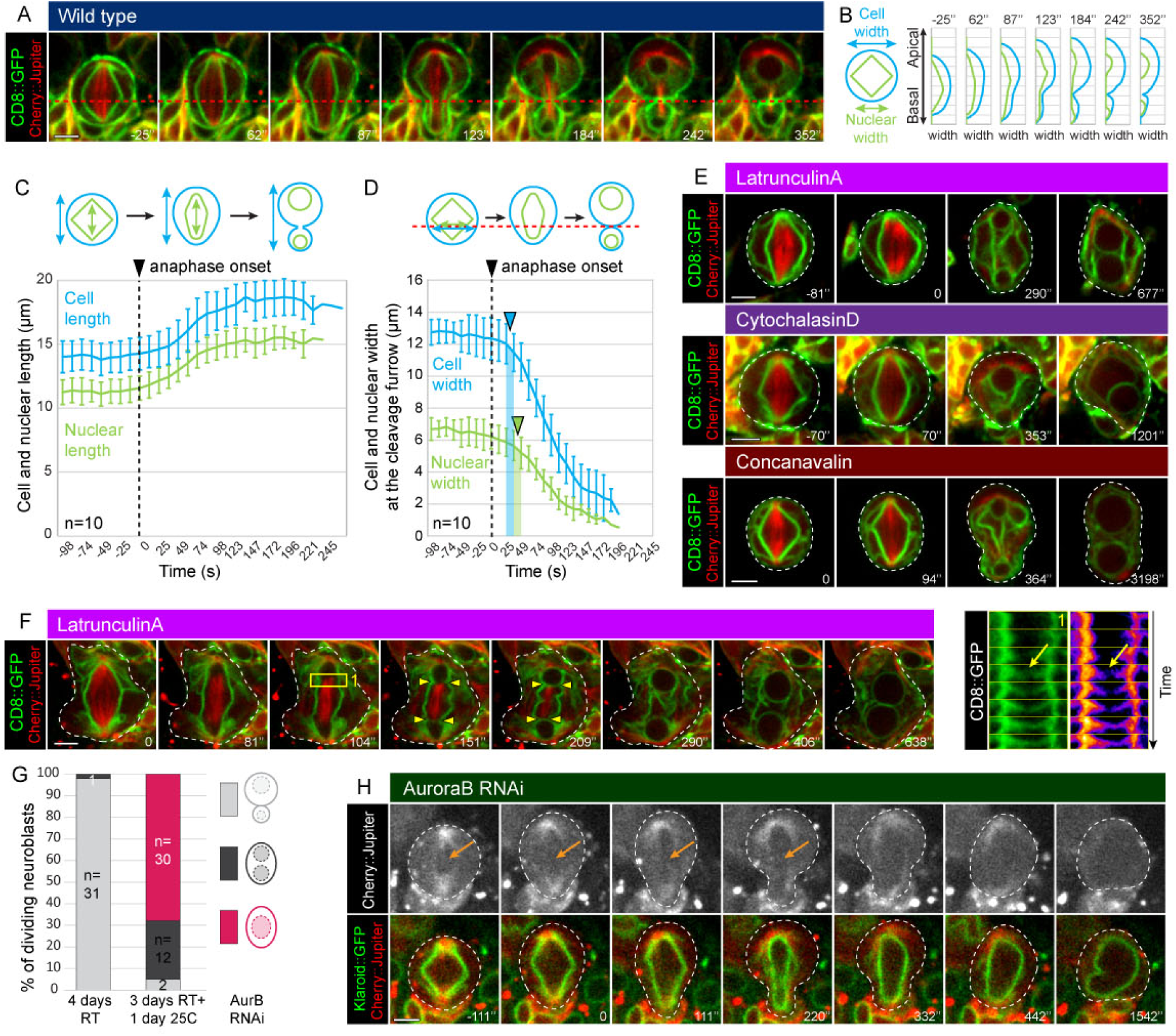
Nuclear division is a sealing-dependent process. (**A**) Dividing neuroblasts imaged in intact brain, expressing the spindle marker Cherry::Jupiter (red) and CD8::GFP (green). The red dashed line represents the site of cortical furrowing. (**B**) Graph showing for one representative cell the cell width (blue curve) and the nuclear one (green curve) from the apical to the basal cortex, for the 7 time points represented in (**A**). (**C**) Graph showing the cell length (blue curve) and the nuclear length (green curve) along the apico-basal axis, averaged from 10 cells, throughout mitosis. Error bars represent standard deviation and the dashed line indicates anaphase onset. (**D**) Graph showing the cell and nuclear width at the cleavage furrow region (along the red dashed line on the cartoon) throughout mitosis, averaged from 10 cells. Error bars represent standard deviation, dashed line indicates anaphase onset, and blue and green arrow heads indicate start of cortical and nuclear furrowing, respectively. (**E**) Representative time lapses of mitotic neuroblasts expressing CD8::GFP (green) and Cherry::Jupiter (red), and failing cytokinesis after LatrunculinA or CytochalasinD treatment or after being cultured on ConcanavalinA. (**F**) Time lapse images of a Latrunculin-treated neuroblast expressing CD8::GFP (green) and Cherry::Jupiter (red). The yellow square represents the area used on the kymograph, and the yellow arrow heads represent the sites of nuclear sealing. (**G**) Quantification of neuroblasts dividing normally (grey category), failing in cell division but still undergoing nuclear division (black category) or failing in both cell and nuclear division (pink category) after expression of an RNAi against AuroraB. The number of cells analysed for each condition is indicated on the graph. (**H**) Representative time lapse images of neuroblast expressing an RNAi against AuroraB, CD8::GFP (green on the merge) and Cherry::Jupiter (white on the top panel, red on the merge), failing in both cell and nuclear division. Orange arrow shows the absence of central spindle. For each experiment, the data were collected from at least 3 independent experiments. Scale bars are 5µm.

These data suggest that it is the central spindle which determines the position of the nuclear sealing sites. Indeed, high temporal resolution images of untreated neuroblasts expressing the INM maker Klaroid::GFP or CD8::GFP revealed sealing events at both extremities of the central spindle (Figure 4A; yellow arrow heads, Figure 4B; yellow arrow heads). The asymmetry of the spindle was translated into asymmetric sealing, as measured by the strong correlation between the nuclear diameter and the centrosome to central spindle distance (Pearson correlation coefficient of 0.9878 (Figure 4C,D)). To further test the idea that it is the asymmetry of the spindle that generates two daughter nuclei with different size as cells leave mitosis, it was important to uncouple spindle asymmetry from the cell division asymmetry. To do so, we overexpressed the polarity protein Galphai. As reported in the literature, in most neuroblasts this led to symmetric divisions with symmetric spindles (Figure S3A) ^19^. Interestingly, however, a subset of neuroblasts divided asymmetrically, despite having near symmetric spindles (Figure 4E; cell division asymmetry ratio of 1.67 and spindle asymmetry ratio of 1.18). Strikingly, nuclear division was close to symmetric in these cells (Figure 4E; nuclear asymmetry ratio of 1.16), as confirmed by a quantitative analysis which showed that the asymmetry in the initial size of nuclei correlated extremely well with the spindle asymmetry at the time of sealing (Figure 4F; left graph; Pearson correlation coefficient r: 0.9893), and did not correlate with the cell division asymmetry (Figure 4F; right graph; Pearson correlation coefficient r: 0.6384). For the converse experiment, to determine whether perturbations that alter asymmetry of the central spindle influence the sites of nuclear sealing and nuclear size, we co-expressed the minus-end microtubule-binding protein Patronin::GFP together with a nanobody fused to the basal localization domain of Pon (GBP-PonLD) and Cherry::Jupiter. As reported in *Drosophila* SOP cells ^20^, the expression of GBP-PonLD was sufficient to recruit Patronin::GFP to the basal cortex of dividing neuroblasts (Figure S3B; yellow arrows). This was associated with an asymmetry in the levels of Patronin::GFP across the central spindle – so that levels were elevated in the basal half of the central spindle rather than apical half (Figure S3B; t300” insert; Figure S3C). Patronin is known to protect the microtubules minus-end from depolymerization by Klp10A ^20-22^ so, as expected, this led to changes in the microtubule-based central spindle. In the control, while the microtubule density was similar on both sides of the central spindle (apical/basal Jupiter intensity ratio: 0.95 ± 0.06) (Figure S3D), the apical half of the central spindle was 1.5 fold longer than the basal half (apical/basal length ratio: 1.54 ± 0.21) (Figure S3E). Basal enrichment of Patronin::GFP led to a local increase in microtubule density and length in this portion of the central spindle, to the detriment of the central spindle in the apical half (apical/basal Jupiter intensity ratio: 0.82 ± 0.09 (Figure S3D) and apical/basal length ratio: 1 ± 0.32 (Figure S3E)). As a consequence, this treatment increased the distance between the apical centrosome and the central spindle and reduced the distance separating the basal centrosome and the central spindle (Figure 4I; left graph). This enhanced both the spindle asymmetry and the nuclear asymmetry (Figure 4G *vs* 4H; green dashed lines, Figure 4I: right graph) without changing the position of the cytokinetic furrow, resulting in an altered cell/nuclear size ratio (Figure S3F). Together, these data emphasize the role of spindle asymmetry inducing an offset that leads to the asymmetric sealing of the nuclear compartment, to generate sibling nuclei with different size.

**Figure 4:**
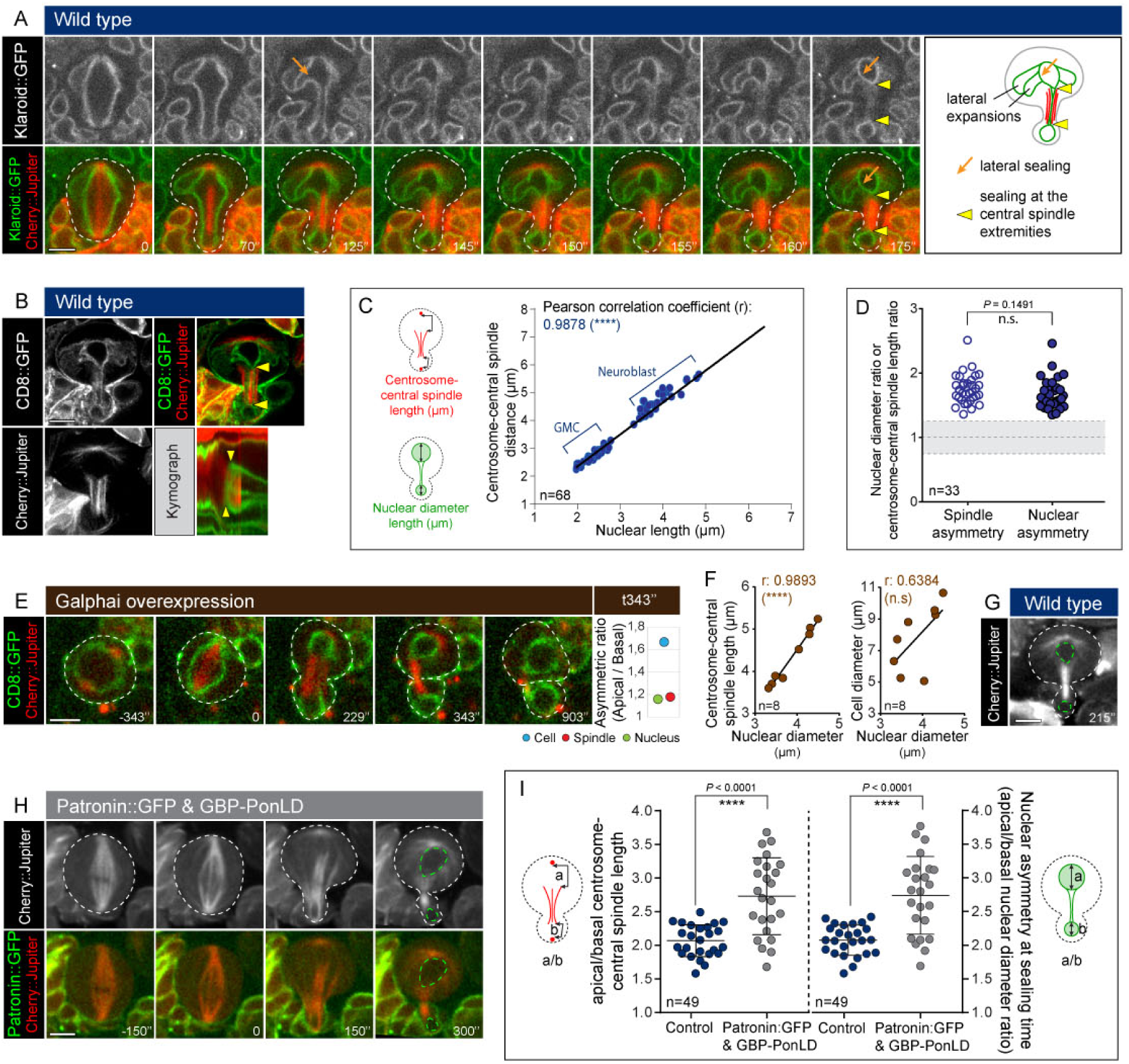
Asymmetric nuclear division depends on nuclear sealing at sites dictated by the spindle. (**A**) Dividing neuroblasts expressing the INM marker Klaroid::GFP (white on top row; green on the merge) and the spindle marker Cherry::Jupiter (red on the merge), imaged with high temporal resolution. Orange arrow shows a lateral envelope sealing event, and yellow arrow heads show sealing sites separating the two daughter nuclei, which are located at both extremities of the central spindle. (**B**) Airyscan pictures of wild type neuroblasts expressing CD8::GFP (white on the top left panel, green on the merge) and Cherry::Jupiter (white on the bottom left panel and red on the merge). Yellow arrow heads show the sealing sites separating the newly formed nuclei, which occur at both extremities of the central spindle. Kymographs are done along the apico-basal axis. (**C**) Plot showing correlation between spindle and nuclear asymmetries. Nuclear length is plotted along the X axis and the distance between the centrosome and the central spindle is plotted along the Y axis, for 68 wild type neuroblasts. (**D**) Graph showing for wild type neuroblasts the spindle asymmetry ratio (empty circles) and the nuclear diameter ratio (filled circles). Number of analysed cells = 33. Asterisks denote statistical significance, derived from unpaired t tests: n.s.: not significant. (**E**) Representative time lapse images of neuroblast expressing CD8::GFP (green), Cherry::Jupiter (red) and Galphai (over-expression). Quantification show the asymmetric ratio for the cell, nucleus and spindle at the sealing time (time point 343s). (**F**) Plot showing correlation between spindle and nuclear asymmetries (on the left) and the absence of correlation between the cell and nuclear asymmetries (on the right). n = 8 for each graph.(**G**) Image of a wild type neuroblast at the nuclear sealing time, showing the symmetry of the central spindle (in term of microtubule density) and the nuclear asymmetry (green dashed line). (**H**) Representative time lapse images of neuroblast expressing Cherry::Jupiter (white on the top panel and red on the merge), Patronin::GFP (green on the merge) and GBP-PonLD (i.e. localization domain of Pon fused to GFP-Binding Protein). Green dashed lines correspond to the nuclei. (**I**) Graph showing the spindle asymmetry (centrosome to central spindle length) (on the left) and the nuclear asymmetry (on the right) for control neuroblasts and for neuroblasts expressing both Patronin::GFP and GBP-PonLD. n = 25 and n = 24 for the control and Patronin::GFP & GBP-PonLD conditions, respectively. Bars indicate mean ± standard deviation. Asterisks denote statistical significance, derived from unpaired t tests: ****: p ≤ 0.0001. For each experiment, the data were collected from at least 3 independent experiments. Scale bars are 5µm.

### Final nuclear sizes are achieved via differential growth of the two daughter nuclei

While the initial asymmetry in nuclear size was established in late anaphase by the sites of sealing at either end of the central spindle, the nuclear asymmetry dramatically increased during telophase and early G1 as the result of differential nuclear growth. Indeed, while the diameter of the neuroblast nucleus was 1.6 larger than that of the GMC nucleus at the time of nuclear sealing (3.89 µm ± 0.39 versus 2.36 µm ± 0.23), it was 3.3 as large by the time nuclear size stabilized ∼10 minutes later (7.70 µm ± 0.43 versus 2.31 µm ± 0.33) (Figure 5A). Since nuclei are roughly spherical, this represents a profound change in the volumetric ratio: the neuroblast nucleus has a volume that is ∼4.5 times larger than the GMC at the time of nuclear sealing, but is ∼37-fold larger by the time nuclear size stabilizes. Quantification of nuclear size over time showed that after nuclear sealing (Figure 5B; step I) neuroblast nuclei continued to grow extensively, while the size of GMC nucleus did not change significantly (Figure 5B; step II), before nuclear size was stabilised (Figure 5B; step III). Importantly, this difference in growth appeared to be related to asymmetries in the distribution of pools of ER membrane required to fuel nuclear envelope expansion. Indeed, we noticed that, at the time of nuclear sealing, multiple lateral sealing events took place in the future neuroblast (Figure 4A; orange arrow, only one lateral sealing event visible at this focal plan) generating a stock of nuclear membrane decorated with ER and INM markers (Figure 5C; yellow arrows). Nuclear growth in the neuroblast was associated with the gradual incorporation of this excess nuclear/ER membrane into the expanding nuclear envelope (Figure 5C; yellow arrows), and halted when the membrane pool was visibly depleted. By contrast, access of the GMC nucleus to this pool of membrane appeared to be restricted by the cytokinetic furrow, preventing its growth.

**Figure 5:**
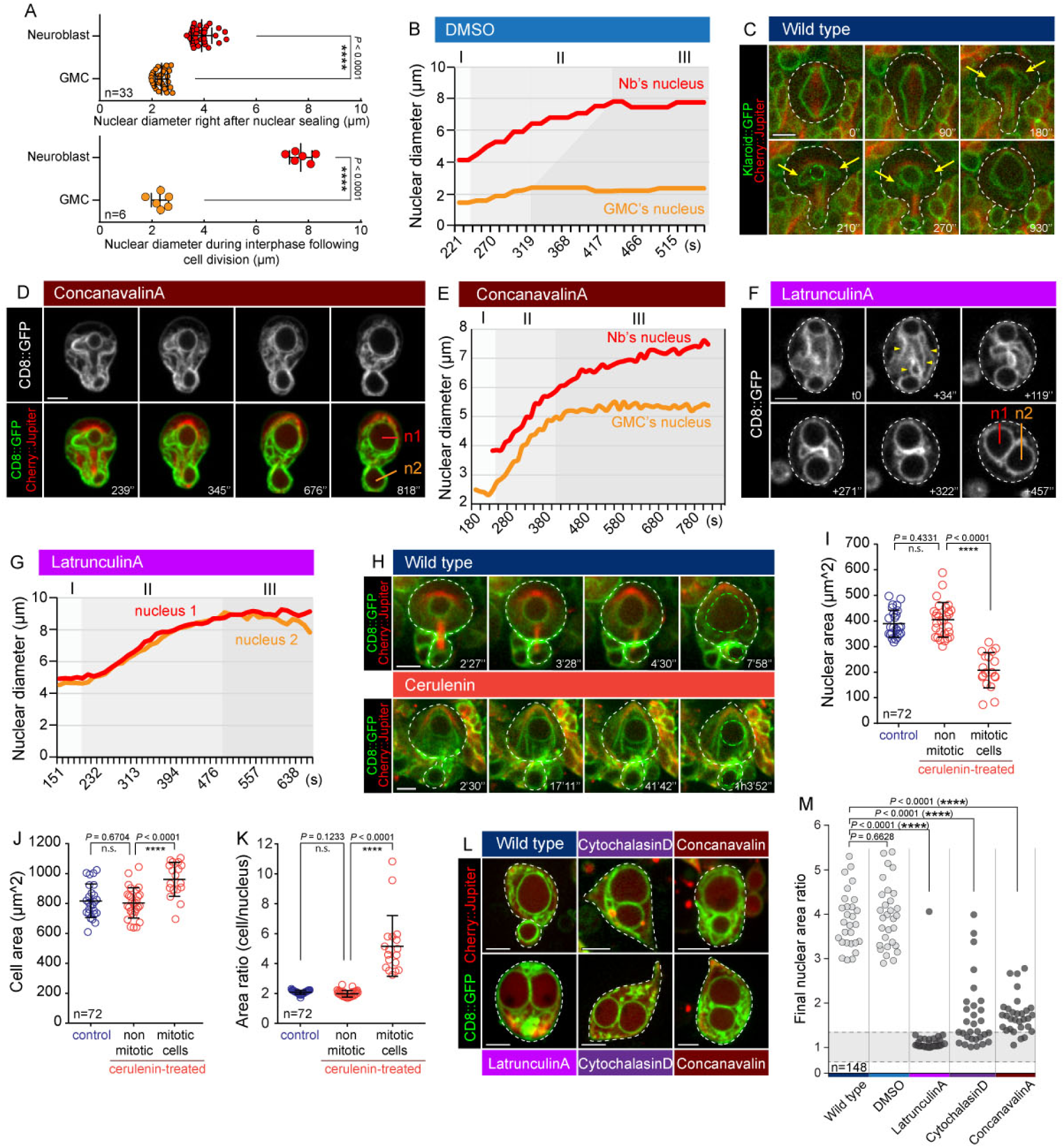
Final nuclear sizes are achieved via differential growth of the two daughter nuclei. (**A**) Scatter plots showing nuclear diameter of neuroblast (red circle) and GMC nucleus (orange circle) right after nuclear formation (top graph) and later during interphase (bottom graph). Bars indicate mean ± standard deviation. Asterisks denote statistical significance, derived from unpaired t tests: ****: p ≤ 0.0001. (**B**) Graph from a representative DMSO-treated neuroblast, showing the nuclear diameter of a neuroblast and GMC nucleus, from nuclei formation to the following interphase. Step I corresponds to the asymmetry establishment, step II to a differential nuclear growth and step III to a plateau. (**C**) Neuroblasts expressing Klaroid::GFP (green) and Cherry::Jupiter (red); yellow arrows show reservoir of nuclear membrane in the cytoplasm of the daughter neuroblast. (**D**) Representative time lapse images of a neuroblast cultured on ConcanavalinA, expressing CD8::GFP (white on the top panel, green on the merge) and Cherry::Jupiter (red on the merge). (**E**) Graph corresponding to the neuroblast presented in panel “d”, showing the nuclear size of both nuclei throughout the cell cycle. Step I corresponds to the nuclei formation, step II to the similar nuclear growth, and step III to the plateau in growth for the GMC nucleus, while the neuroblast continues to grow. (**F**) Representative time lapse images of a Latrunculin-treated neuroblast expressing CD8::GFP (white). Yellow arrow heads indicate the reservoir of nuclear membrane available in the cytoplasm, which progressively disappears as nuclei grow. (**G**) Graph from a representative Latrunculin-treated neuroblast showing the size of both nuclei throughout the cell cycle. Step I corresponds to nuclear formation, step II to nuclear growth and step III to the plateau representing the end of growth. (**H**) Untreated (top panel) and Cerulenin-treated neuroblast (bottom panel) expressing CD8::GFP (green) and Cherry::Jupiter (red). (**I**) Graph showing nuclear area for control neuroblasts and Cerulenin-treated neuroblasts (non-mitotic and mitotic). n = 72. Bars indicate mean ± standard deviation. Asterisks denote statistical significance, derived from unpaired t tests: n.s.: not significant, ****: p ≤ 0.0001. (**J**) Graph showing cell area for control neuroblasts and Cerulenin-treated neuroblasts (non-mitotic and mitotic). n = 72. Bars indicate mean ± standard deviation. Asterisks denote statistical significance, derived from unpaired t tests: n.s.: not significant, ****: p ≤ 0.0001. (**K**) Graph showing area ratio (cell/nucleus) for control neuroblasts and Cerulenin-treated neuroblasts (non-mitotic and mitotic). n = 72. Bars indicate mean ± standard deviation. Asterisks denote statistical significance, derived from unpaired t tests: n.s.: not significant, ****: p ≤ 0.0001. (**L**) Representative pictures showing wild type and binucleated neuroblasts expressing CD8::GFP (green) and Cherry::Jupiter (red), containing 2 nuclei with either different or similar size. (**M**) Scatterplot showing the nuclear area ratio between the neuroblast and the GMC nuclei for both wild type and DMSO conditions (grey circles), or between the two daughter nuclei of binucleated cells after Latrunculin A, Cytochalasin D or Concanavalin A treatment (black circles). Grey area on the graph represents a nuclear area ratio equal to 1 ± 0.25. Number of analysed cells = 148. Asterisks denote statistical significance, derived from unpaired t tests: n.s.: not significant, ****: p ≤ 0.0001. For each experiment, the data were collected from at least 3 independent experiments. Scale bars are 5µm.

To test if the constriction induced by the cytokinesis furrow plays a role in limiting the availability of the total ER pool available to neuroblast and GMC nuclei as they grow, as suggested by these data, we used treatments to induce division failures at different stages in the process of cytokinesis. In cells plated on Concanavalin A-coated dishes, where furrow contraction is slowed by adhesion to the dish ^17^, nuclei began to grow at similar rates, being in contact with the same ER pool (Figure 5D; time points 239” and 345”, Figure 5E; step II). However, once the furrow was sufficiently narrow to block access to the bulk of cytoplasmic ER, GMC nuclear growth stopped, whilst the neuroblast nucleus continued to expand (Figure 5D; time point 818”, Figure 5E; step III). As a consequence, when cytokinesis failed at very late stages, this yielded single cells containing one large and one small nucleus. However, when we used the actin-poison Latrunculin A to block both the formation and contraction of the cleavage furrow, both nuclei displayed a similar growth rate (Figure 5F,G; step II) before reaching a plateau at the same time (step III), yielding two nuclei of identical size (Figure 5F). Note that since Latrunculin treatment perturbs cortical polarity ^23^, an important regulator of spindle asymmetry ^24, 25^, neuroblasts treated with Latrunculin enter anaphase with a symmetrical spindle (Figure 3F), leading to the formation of sibling nuclei with identical size (Figure 5F, G; step I).

If nuclear growth depends on the ER membrane reservoir, as suggested by this analysis, treatments that interfere with lipid synthesis should reduce nuclear size. To test whether or not this is the case, we treated neuroblasts with Cerulenin, an inhibitor of B-keto-acyl-ACP synthase and HMG-CoA synthase involved in fatty acid and sterol biosynthesis ^26, 27^. This treatment impaired the growth of the neuroblast nucleus (Figure 5H; bottom panel). Importantly, this was only the case for neuroblasts that were already in mitosis at the time of drug addition (Figure 5I), implying that the reduction of the nuclear growth is not an indirect consequence of lipid metabolism deregulation during interphase. Furthermore, this was not associated with a reduction in cell size (Figure 5J). As a result, we observed an increase in the nuclear / cell size ratio in Cerulenin-treated mitotic neuroblasts (Figure 5K). This makes it clear that fatty acids/sterol synthesis and membrane addition are limiting factors for nuclear growth. Thus, nuclear growth is not a simple consequence of the movement of cytoplasmic material into the nucleus, as has been suggested in other model systems ^28-30^.

Taken together, these data show that differences in the final size of sibling nuclei depend on two processes that occur in sequence: i) asymmetric nuclear sealing controlled by the spindle geometry, and ii) differential nuclear growth which depends on local ER availability. Thus, depending on spindle asymmetry, cleavage furrow dynamics and the timing of cytokinesis failure, binucleate cells can have nuclei that are similar or very different in size (Figure 4L, M).

### Asymmetries in the nuclear and chromatin organization in neuroblast divisions

In the wildtype, this multi-step process of asymmetric nuclear division generates two nuclei that differ in their molecular composition (e.g. nucleoporins) (Figure 1D, Figure S1E) and size (Figure 4, 5), despite their both carrying the same amount of DNA. This led to asymmetries in nuclear packing. To visualise when during mitotic exit this change in packing occurs, we analysed cells leaving mitosis using the DNA marker Histone 2A::mRFP to mark chromatin, together with the cortical protein Spaghetti Squash::GFP. The two sets of sister chromatids appeared identical immediately after DNA segregation at anaphase (Figure 6A; time points 30”-120”). However, as cells progressed through telophase into interphase, the His2A signal was seen becoming weaker in the neuroblast nucleus, relative to the GMC nucleus (Figure 6A; time point 840”, 6B), as the DNA spread to fill the entire nuclear compartment (Figure S4A). A similar effect was seen using DAPI staining (Figure S4B). We also examined the same process in neuroblasts that fail in cytokinesis. As expected, the DAPI signal between sibling nuclei was very different in binucleated neuroblasts displaying two nuclei with different size (Figure S4C; left graph), but was similar when both nuclei display identical size (Figure S4C; right graph). We also stained neuroblasts for H3K4me2, a Histone post-translational modification that has been associated with an increase of transcriptional activity in various model systems including in *Drosophila* ^*31, 32*^, which increases during differentiation of neural stem cells into neural progenitors in mouse brain ^33^. This revealed a stronger net H3K4me2 signal in small nuclei (Figure 6C; yellow arrow), relative to the DAPI control (Figure 6D; DAPI sum intensity ratio of 1.074 ± 0.12 versus H3K4me2 sum intensity ratio of 0.77 ± 0.09). In fact, in untreated dividing neuroblasts, H3K4me2 staining of segregating chromatids appeared symmetric until nuclear sealing in late anaphase (Figure 6E; yellow arrows), after which it became more intense in small nuclei relative to large neuroblast nuclei (Figure S4D; yellow dashed lines and yellow stars). Thus, the physical asymmetry of the nuclear division is associated with asymmetries in chromatin organization, which include H3K4me2 - a hallmark of neural differentiation ^33^.

**Figure 6:**
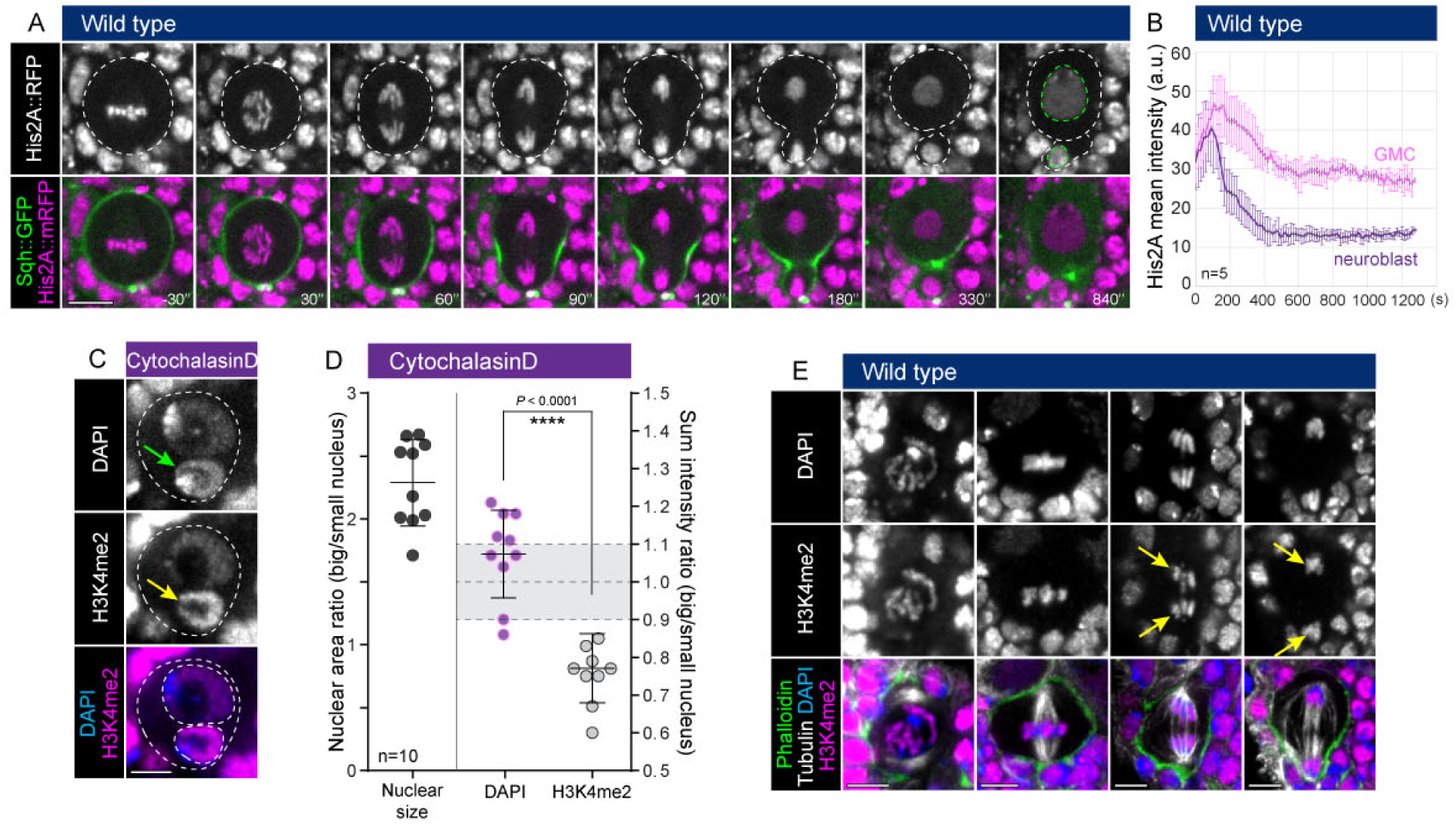
Asymmetries in the nuclear and chromatin organization in neuroblast divisions. (**A**) Representative time lapse images of a neuroblast expressing the DNA marker His2A::RFP (white on the top panel and pink on the merge) and the cortical marker Sqh::GFP (green on the merge). (**B**) Graph showing His2A::RFP mean intensity over time, into the GMC (pink curve) and neuroblast nucleus (purple curve). n = 5. Error bars represent standard deviation. (**C**) Binucleated neuroblast (Cytochalasin-treated) fixed and stained for DAPI (white on the top panel and blue on the merge) and H3K4me2 (white on the middle panel and pink on the merge). Green and yellow arrows show the higher intensity of DAPI and H3K4me2 in the small nucleus. (**D**) Graph showing the nuclear area ratio (on the left) and the DAPI or H3K4me2 sum intensity ratio (on the right) in binucleated neuroblasts displaying two nuclei with different size. n = 10. Bars indicate mean ± standard deviation. Asterisks denote statistical significance, derived from unpaired t tests: ****: p ≤ 0.0001. (**E**) Wild type neuroblast fixed and stained for DAPI (white on the top panel and blue on the merge), H3K4me2 (white on the middle panel and pink on the merge), Tubulin (white on the merge) and Phalloidin (green on the merge). Yellow arrows show the similar intensity of H3K4me2 on the two pools of sister chromatids. For each experiment, the data were collected from at least 3 independent experiments. Scale bars are 5µm.

### Nuclear identity is set by the relative timing of nuclear division, cell division and cortical release of nuclear proteins

Finally, having shown that the nuclear division contributes to the formation of two nuclei of different size and composition, and knowing that the cortical polarity is important to generate two cells of different fate, we wanted to determine how nuclear and cell division are coordinated in time and space to generate two nuclei with different identities. For this analysis, we wanted a system in which to compare nuclear size and composition in cases of early versus late cell division failure. To do so, we treated neuroblasts expressing Pebble::GFP (Ect2 ortholog) with Cytochalasin to induce a late division failure - using Latrunculin as a control to induce an early cytokinesis failure. In these experiments, Pebble served as an excellent marker of cortical polarity, mitotic progression and nuclear import (Figure S5A). In absence of treatment, Pebble::GFP was asymmetrically localized at the cortex during mitosis, and was 1.5 (± 0.3) fold enriched in the neuroblast nucleus relative to that of the GMC after its nuclear import by late telophase (Figure 7A,B and Supplemental Video 7). While Pebble was still asymmetrically recruited to the cortex in both Latrunculin- or Cytochalasin-treated neuroblasts (Figure S5B,C), its differential nuclear accumulation was lost in cells failing early in cytokinesis (Latrunculin treatment; Figure 7B,C and Supplemental Video 8). By contrast, cells undergoing a late cytokinesis failure formed two nuclei of different sizes that contained different concentrations of Pebble (Cytochalasin treatment; Figure 7B, d and Supplemental Video 9), which were indistinguishable from those in untreated neuroblasts (Figure 7B). This striking result led us to examine whether the same might be true of endogenous cell fate determinants that are, like Pebble::GFP, asymmetrically localised along the cortex before being released and imported into nuclei during telophase. To do so, we analysed the distribution of Prospero, a transcription factor that acts as a switch between self-renewal and differentiation in neural stem cells ^34^. As previously reported, in control cells Prospero was found sequestered at the basal cortex as the result of its interaction with the polarity protein Miranda ^35^, before being released and imported into the GMC nucleus after cytokinesis (Figure 5D; white arrows). In cells undergoing a very late cytokinesis failure (Cytochalasin treatment in combination with Pebble overexpression), the closure of the cleavage furrow prevented Prospero from diffusing through the cytokinetic bridge after its release from the basal cortex. As a result, it only entered the smaller GMC nucleus. However, following abscission failure in late telophase/interphase, the two daughter nuclei containing different levels of the cell fate marker Prospero entered the same cytoplasm (Figure 7E). Thus, while both physical and molecular nuclear asymmetries were lost from cells that underwent an early cytokinesis failure, late cytokinesis failures generated cells with two nuclei that differ profoundly in their size, composition and complement of cell fate determinants despite being situated in the same cytoplasm. Moreover, these differences were found to persist with time (Figure 7F,G & Figure S5E), implying that nuclear size and composition are fixed once established in early G1. Hence, our study shows that the polarization of cell fate determinants *per se* is not sufficient to generate two daughter nuclei with different identities. It is the relative timing of (i) nuclear division, (ii) polarity-dependent cortical release of nuclear proteins and (iii) cytokinesis that serves to generate two daughter nuclei with different identities (i.e. size and composition including transcription factors) and, as a consequence, different fates.

**Figure 7:**
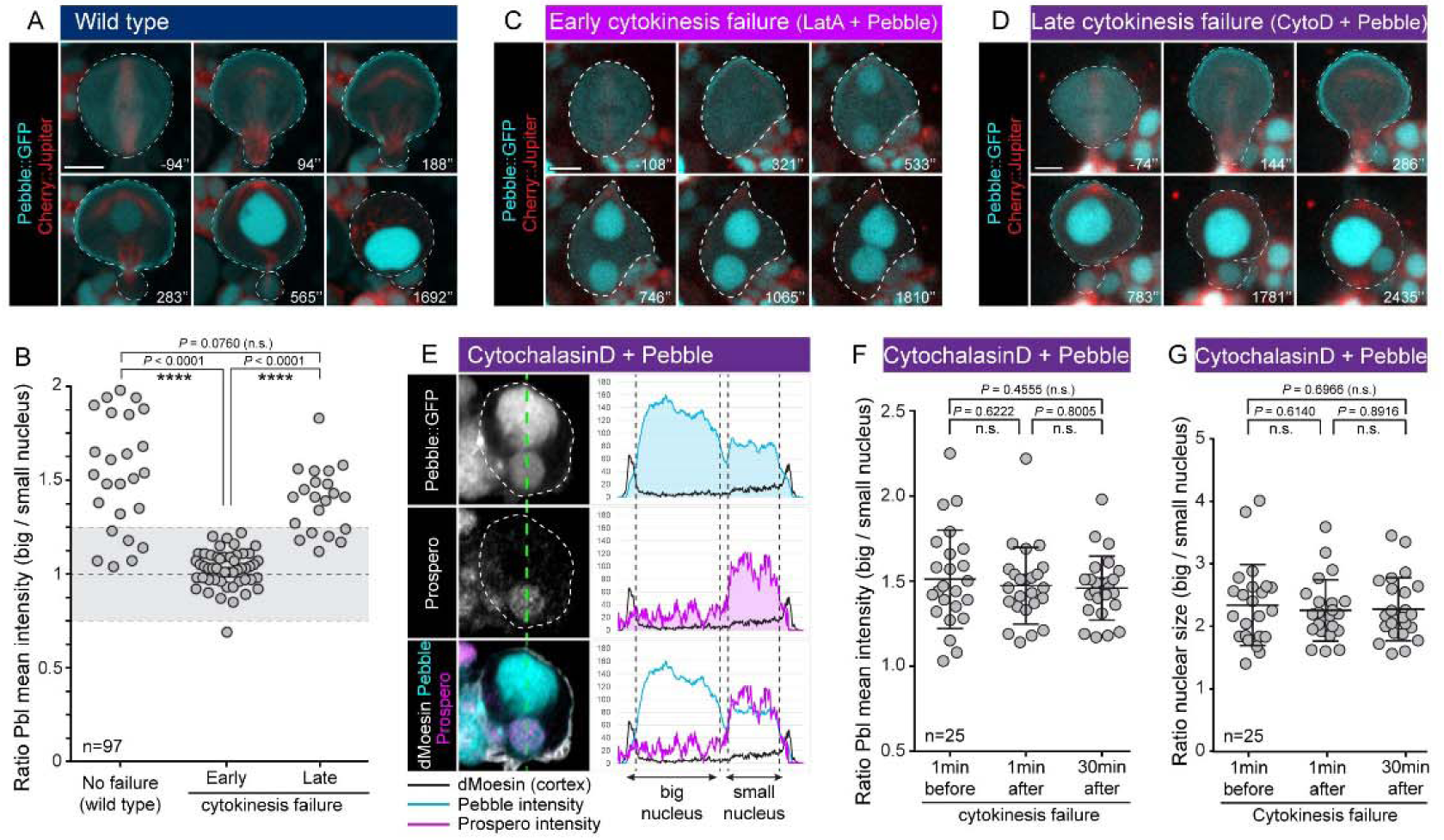
Nuclear identity is set by the relative timing of nuclear division, cell division and cortical release of nuclear proteins. (**A**) Representative neuroblasts expressing Pebble::GFP (blue) and Cherry::Jupiter (red). (**B**) Scatterplot showing the Pebble::GFP mean intensity ratio of the two daughter nuclei in neuroblasts dividing normally, failing early in cytokinesis or failing late. Number of analysed cells = 97. Asterisks denote statistical significance, derived from unpaired t tests: n.s.: not significant, ****: p ≤ 0.0001. (**C, D**) Representative neuroblast expressing Pebble::GFP (blue) and Cherry::Jupiter (red) failing (**C**) early in cytokinesis after LatrunculinA treatment or (**D**) late during telophase/interphase after Cytochalasin treatment. (**E**) Immunostaining of binucleated neuroblasts expressing Pebble::GFP (white on the top panel; blue on the merge) and treated with Cytochalasin, for the transcription factor Prospero (white on the middle panel; pink on the merge) and the cortical marker dMoesin (white on the merge). Graphs show the intensity of the different markers along the green dashed line going through the middle of both nuclei. (**F**) Scatterplot showing the Pebble::GFP mean intensity ratio of the two nuclei in binucleated Cytochalasin-treated neuroblasts over-expressing Pebble::GFP, 1 minute before or after cytokinesis failure and 30 minutes later. Number of analysed cells = 25. Bars indicate mean ± standard deviation. Asterisks denote statistical significance derived from unpaired t tests: n.s.: not significant. (**G**) Scatterplot showing the nuclear size ratio between the two nuclei of binucleated Cytochalasin-treated neuroblasts over-expressing Pebble::GFP, 1 minute before, or after cytokinesis failure, and 30 minutes later. Number of analysed cells = 25. Bars indicate mean ± standard deviation. Asterisks denote statistical significance, derived from unpaired t tests: n.s.: not significant. For each experiment, the data were collected from at least 3 independent experiments. Scale bars are 5µm.

## Discussion

### A mitotic nuclear compartment

Here, through the analysis of the nuclear compartment during asymmetric neuroblast divisions, we show that nuclei are not dissolved and reformed during passage through mitosis as they are in mammalian cell divisions. Instead, the nuclear envelope in the fly neuroblast is remodelled so that it persists throughout mitosis as a membrane that contains INM and ONM markers (SUN/KASH proteins, Klaroid and Klarsicht respectively), together with a supporting underlying lamina which envelopes the spindle. This is the case even though the nuclear envelope loses nuclear pore complexes and its barrier function upon mitotic entry. As revealed by electron microscopy, this is because the mitotic nuclear membrane does not form a tight and continuous barrier, but rather exists as a discontinuous multi-layered structure that appears to be sufficiently well organised to keep large structures (such as vesicles and mitochondria) out. While others have evoked a similar function for a putative spindle matrix ^36^, the persistence of a nuclear compartment may also enable the nuclear membrane to be inherited rather than being lost and re-established *de novo* – aiding rapid nuclear reformation and asymmetric nuclear formation. These differences between mitosis in fly neural stem cells and mammalian cells appear to be at least in part due to the persistence of a lamina in mitotic fly neuroblasts ^16^. While the fragmentation of the mitotic neuroblast nuclear envelope could be induced using Lamin RNAi, these cells were unable to reassemble a nucleus as they re-entered G1; implying that *Drosophila* neuroblasts likely require the maintenance of this nuclear envelope for efficient nuclear formation at mitotic exit and that they cannot rely on the processes thus far described for human cells ^37^.

### Forming two nuclei of different size

Importantly, while this study also demonstrates that nuclear division of *Drosophila* neuroblasts is an autonomous mechanism that is independent of cytokinesis, it is clear from our analysis that the asymmetry in nuclear division depends on factors extrinsic to the chromatin and nuclear envelope. Thus, asymmetric division of the nuclear compartment to generate one small and one large nucleus appears to depend on pre-existing asymmetries in the cortex and spindle. This is achieved in two steps. First, the central spindle defines the sites where the nuclear compartment is sealed. This separates the nuclear membrane into three pools, and generates two physically distinct nascent daughter nuclei with different size. While further investigation is required to determine the molecular mechanism of nuclear resealing, ESCRTIII proteins are likely to play a key role, based on their conserved role in repairing holes in the nuclear membrane in yeast and mammalian cells exiting mitosis ^38, 39^.

Second, these nuclei compete for the available reservoir of ER and nuclear membrane as they grow. Importantly, access to this membrane reservoir appears to be limited by partial closure of the cytokinesis furrow, since pinching of the plasma membrane appeared to be sufficient to limit nuclear growth before abscission. This suggests the presence of a barrier at the neck of the cytokinesis furrow in fly neuroblasts that functions like the neck of dividing budding yeast cells ^40, 41^, which is able to restrict the movement of material between the two asymmetrically daughter cells prior to division. However, we were unable to identify a clear role for Septins in this process (data not shown), as was observed in yeast ^40, 41^.

### Functional implications of asymmetric nuclear division

Physical asymmetries in the two daughter nuclei generated by nuclear division are likely to have profound functional consequences. This is suggested by our data which show that, asymmetries in nuclei sealing and in nuclear growth yield a 37-fold difference in volume between sibling nuclei. This size asymmetry had a strong impact on chromatin packing and is associated with a differential accumulation of H3K4me2, a hallmark of actively transcribed genes involved in neural differentiation, in small versus large nuclei. We also found that the concentration of NLS::GFP remained the same irrespective of nuclear size (data not shown). This suggests an equivalent asymmetry in the number of molecules imported from the cytoplasm into each nucleus. These molecular asymmetries are then likely to be further accentuated by differences in the composition of the nuclear envelope in the two nuclei, e.g. NPCs and the lamina. These were readily apparent in our images of neural stem cell divisions (Figure 1D, 2A; yellow arrows, Figure S1E). Remarkably, this asymmetry in the structure and size of the sibling nuclei occured long before cortical cell fate release and cytokinesis. For this reason, the asymmetry of the nuclear division likely helps generate sibling nuclei with different identities to generate daughter cells of different fate. Thus, the impact of nuclear asymmetries upon entry into G1 on nascent transcription is something that now requires much further exploration.

It is also clear from our analysis that the relative timing of nuclear and cell division plays an important role in generating functionally asymmetric divisions. As we show here, nuclei are formed prior to the release of cortically-localised transcription factors. In order to generate two cells with different fates, cytokinesis must occur before cortical release. As a result, a late division failure yielded daughter nuclei that differed in the levels of asymmetrically localised cortical factors, such as Pebble::GFP and the cell fate determinant Prospero. One might ask why the system is set up in this way. While further work is required to answer this question, our analysis hints at this being a way to rapidly define nuclear identify very early in G1, which might be particularly important in these cells due to their very short cell cycle (45 min to 1 hour on average). It is also possible that this system allows nascent nuclei to compete for common factors prior to cell division, facilitating the break symmetry required to distinguish the neural stem cell from the GMC (Figure 1). Finally, we believe this first detailed analysis of the nuclear division process in fly neural stem cells will open up new avenues of research into the mechanism via which nuclear division contributes to the acquisition of distinct cell fates.

## Acknowledgments

We would like to thank Andrew Vaughan and John Gallagher for microscopy support; Talila Volk, Saverio Brogna, Christian Lehner, François Payre and Emmanuel Derivery for reagents; as well as Franck Pichaud, Yanlan Mao, Jens Januschke, Helen Matthews, André Pulschen, Emmanuel Derivery and the Baum lab for feedback on this manuscript. This work was supported by the MRC LMCB, a Cancer Research UK programme grant (C1529/A17343) and a BBSRC project grant (BB/R009732/1).

## Contributions

CR and BB conceived the project. CR performed and analysed the experiments except the electron microscopy. IJW performed the electron microscopy experiments. CR and BB wrote the manuscript.

## Materials and methods

### Experimental model

Several lines of the fruit fly Drosophila melanogaster were used in this study. Flies and larvae were raised in vials containing standard cornmeal-agar medium supplemented with baker’s yeast and incubated at 25°C. Third-instar larvae of 3-days old (expressing live imaging markers) or 5-days old (expressing RNAi) were dissected to extract the brains, then live imaging was performed directly on brains or after brain dissociation (see “Method details”). Experiments utilized both female and male larvae, and data from both sexes was pooled.

### Fly genetics

Flies used in this study include: Klaroid^CB04483^ (BDSC-51525) ^42^, UAS BiP::sfGFP::HDEL (BDSC-64749) ^43^, UAS Klarsicht::GFP ^44^, UAS RpS13::GFP ^45^, gNup58::EGFP ^46^, EGFP::Nup107 (BDSC-35514) ^10^, UAS LamDm0::GFP (BDSC-7376), UAS 5EGFP::NLS (BDSC-65402), UAS Pebble::GFP ^47^, UAS dsRNA LamDm0 (BDSC-36617 and BDSC-57501), UAS Cherry::Jupiter, WorGal4, UAS Cherry::Jupiter; UAS CD8::GFP (this study), WorGal4, Klaroid^CB04483^, UAS Cherry::Jupiter; UAS Dicer (this study), UAS Galphai (BDSC-44600), w; UAS GBP::PonLD; UAS GFP::Patronin^20^, UAS AuroraB RNAi (VDRC-35107) ^48^, UAS AuroraB RNAi (BDSC-35299)^48^, w; WorGal4, UAS His2A::mRFP ^49^, Sqh::GFP ^50^. Transgenes were expressed using the neuroblast-specific driver worGal4 ^24^.

### Live imaging sample preparation

For experiments performed on intact brains, imaging medium (Schneider’s insect medium mixed with 10% FBS (Sigma), 2% PenStrepNeo (Sigma), 0.02 mg/mL insulin (Sigma), 20mM L-glutamine (Sigma), 0.04 mg/mL L-glutathione (Sigma) and 5 µg/mL 20-hydroxyecdysone (Sigma)) was warmed up to room temperature before use. Seventy-two hours after egg laying, larvae were dissected in imaging medium then brains were transferred onto a gas-permeable membrane (YSI Life Sciences #13-298-83) fitted on a metallic slide. Brains were oriented with the brain lobes facing the coverslip, and excess media was removed until the brain lobes were in contact with the coverslip. The sample was sealed with Vaseline.

### Primary neuroblast cultures

For experiments done on primary neuroblast culture, seventy-two hours larvae were dissected in Chang & Gerhing solution (3.2 g/L NaCl, 3 g/L KCL, 0.69 g/L CaCl2-2H2O, 3.7 g/L MgSO4-7H2O, 1.79 g/L tricine buffer pH 7, 3.6 g/L glucose, 17.1 g/L sucrose, 1 g/L BSA) ^51^ at room temperature. Brains were then incubated in Chang & Gerhing solution supplemented in collagenase from Clostridium histolyticum (Sigma) and papain from papaya latex (Sigma) at a final concentration of 1 mg/mL each, during 45 minutes at 29 °C. Brains were washed with imaging medium (see above) then dissociated in imaging medium by pipetting 20–25 times. The cell culture was then imaged using Ibidi chambers (15µ-slide angiogenesis).

### Antibodies and immunostaining

The following primary antibodies were used for this study: mouse anti-LamDm0 Ser25 (DHSB; #ADL84.12; 1:100), mouse anti-LamDm0 (DHSB; #ADL67.10; 1:100), chicken anti-GFP (Abcam; #ab13970; 1:1000), rabbit anti-dMoesin (gift from Francois Payre; 1:1000), mouse anti-Prospero (DHSB; #MR1A; 1:100), rabbit anti-H3K4me2 (Abcam; #ab7766; 1/100), and rabbit anti-Miranda (gift from Emmanuel Caussinus; 1:50). Secondary antibodies were from Invitrogen and Phalloidin-FITC (Sigma; #P5282; 1:100) was used to stain F-Actin. For immunostaining, seventy-two hours larvae were dissected in Schneider’s insect medium (Sigma-Aldrich S0146) and brains were fixed for 20 min in 4% paraformaldehyde in PEM (100 mM PIPES pH 6.9, 1 mM EGTA and 1 mM MgSO4). After fixing, the brains were washed with PBSBT (1× PBS (pH7,4), 0.1% Triton-X-100 and 1% BSA) then blocked with PBSBT for 1 h. Primary antibody dilution was prepared in PBSBT and brains were incubated 48 h at 4 °C. Brains were washed with PBSBT four times for 30 min each, then incubated with secondary antibodies diluted in PBSBT at 4 °C overnight. The next day, brains were washed with PBST (1x PBS, 0.1% Triton-X-100) four times for 20 min each and kept in Vectashield antifade mounting medium with DAPI (Vector laboratories; H1200), at 4 °C.

### Small molecule inhibitors and chemical treatments

Following dissection and prior montage for live imaging, brains were incubated in imaging medium (see above) containing the following chemical reagents, as indicated on the Figures: Latrunculin A (Sigma; #L5163; final concentration of 10 µM), Cytochalasin D (Sigma; #C8273; final concentration of 5 µg/mL or 10 µg/mL) or Binuclein 2 (Sigma; #B1186; final concentration of 10 µM or 20 µM). Cerulenin (Sigma; #C2389; 10µM) was added during live imaging. After brain dissociation, the following final concentrations were used on primary neuroblast culture: Latrunculin A: 1 µM; Cytochalasin D: 0.5 µg/mL. For control experiments, dissected brains were incubated in DMSO at the same final concentrations than the ones used to dissolve the reagents. In order to induce binucleated cells, primary neuroblast culture was platted on dish coated with Concanavalin A (Sigma # L7647). Observation of mitochondria distribution during mitosis was done by incubating primary neuroblast culture with MitoTracker Deep Red (Invitrogen # M22426; final concentration of 0.1nM).

### Electron microscopy

Cells were cultured on gridded coverslip-bottomed dishes (MatTek, P35G-1.5–14-CGRD) to facilitate correlation between light and electron microscopy, and processed for the latter following a protocol adapted from Deerinck *et al* (NCMIR methods for 3D EM: A new protocol for preparation of biological specimens for serial block face scanning electron microscopy). Briefly, coverslips were fixed in 2% PFA / 2.5% Gluteraldehyde solution (EM grade, TAAB) in 0.1M sodium cacodylate buffer for 30 min at room temperature, washed in 0.1M sodium cacodylate buffer and post-fixed in 1% OsO_4_/1.5% potassium ferricyanide for 1 hour at 4°C. Samples were then treated with 1% thiocarbohydrazide (TCH) for 20 min at room temperature, 2% OsO_4_ for 30 min at room temperature, 1% uranyl acetate (UA) overnight at 4°C and lead aspartate for 30 min at 60°C, with washing in dH_2_O between each step. This was followed by dehydration of the samples by graded ethanol incubations in 70%, 90% and 100 % ethanol, and incubations for 1 hour in a 1:1 mix of propylene oxide: epon resin (TAAB), 1 hour in fresh 100% epon resin, and a change to a final 1 hour epon resin incubation all at room temperature before mounting the coverslip by inversion onto a pre-polymerised epon resin stub and polymerisation overnight at 60°C. Coverslips were removed by plunging into liquid nitrogen. Cells of interest were relocated on the blockface using the transferred alphanumeric grid with reference to light microscopy images, and the resin block trimmed accordingly. Ultrathin sections of 70 nm were cut using a diamond ultra 45-degree knife (Diatome) on a Leica UC7 ultramicrotome, and collected on formvar coated copper 2 x 1mm slot grids. EM images were acquired on a transmission electron microscope (T12 Tecnai Spirit biotwin, FEI).

### Confocal microscopy

Fixed samples were imaged using an inverted Leica SP5 confocal microscope. For representative images, a 60×/1.40 N.A oil immersion objective was used. Live cell imaging was performed on a UltraView Vox spinning disc confocal microscope (Perkin Elmer Nikon TiE; Yokogawa CSU-X1 spinning disc scan head) with 60×/1.40 N.A oil objective and equipped with a Hamamatsu C9100-13 EMCCD camera, or a 3I spinning disc confocal microscope (Zeiss AxioObserver Z1; Yokogawa CSU-W1 spinning disc scan head) with 63×/1.40 N.A objective and equipped with a photometrics prime 95B scientific CMOS camera. Both spinning disc microscopes are equipped with a temperature-controlled environment chamber set at 26C for the experiments.

### Image processing and calculations

Images were processed using Imaris × 64 7.5.2 and ImageJ softwares. Measurements of intensity was obtained by using the oblique slicer in Imaris oriented along the cell division axis and going through the mid-plan of both daughter nuclei (neuroblast and GMC, respectively). The corresponding images were exported as TIFF files and opened with ImageJ to measure the mean intensity or integrated density, as mentioned on graphs. To measure the ratio cytoplasmic / nuclear NLS::5GFP or Rps13::GFP, background correction was performed by measuring the green channel intensity in the media. For the cell width and nuclear width values over time, a grid was used to measure the cell and nuclear width along each of the line of the grid, starting from the apical cortex towards the basal one for each time point. Cell and nuclear furrowing values were obtained by measuring over time both cell and nuclear width along the red line, which represents the cleavage furrow plan. To do so, images were exported as TIFF then combined in a stack using ImageJ. The red line was drawn at anaphase when the cleavage furrow positioning is set and easy to determine. Measurements of both compartment widths were then done along this line, prior and after anaphase onset, to quantify the cleavage furrow ingression and the deformation of the nuclear compartment at the cleavage furrow region. Measurements of nuclear or cell area ratio were performed by using oblique slicer in Imaris as previously described, then images were exported as TIFF and opened with ImageJ to measure manually the area or perimeter (as mentioned on the figures) by drawing the border of each nucleus and cell. For the graphs showing nuclear growth, TIFF files were opened with ImageJ then the nuclear diameter was manually measured for each time point. Microtubule density was based on the measurement of Cherry::Jupiter intensity, as reported in *Derivery & al*^*20*^. To measure the sum intensity of DAPI and H3K4me2, maximal projection was done using Imaris then exported into ImageJ. Measurements were performed with background subtraction. For the graph showing Pebble distribution in untreated, Latrunculin A- and Cytochalasin D-treated neuroblasts, a line was drawn along the cortex starting from the apical region for three different time points, then the profiles obtained were plotted on radar graphs. For the plot profiles of Pebble, Prospero and dMoesin in binucleated cells, a line was drawn going through the middle of both nuclei then corresponding plot profiles were obtained using ImageJ and values were exported on Excel. All figures were assembled using Adobe Illustrator CS6 software.

### Statistical analysis

For each experiment, the data was collected from at least 3 independent experiments. For each independent experiment, at least 8 larvae were dissected. For all the analysis, “n” refers to the number of cells analysed and is mentioned on each graph as well as in figure legends. Statistical significance was determined with Student’s t test using GraphPad Prism software. In all Figures the Prism convention is used: n.s. (P > 0.05), *(P ≤ 0.05), **(P ≤ 0.01), ***(P ≤ 0.001) and ****(P ≤ 0.0001). In all graphs showing mean, the error bars correspond to standard deviation.

### Data and code availability

This study did not generate code.

## Supplemental information

**Figure S1:**
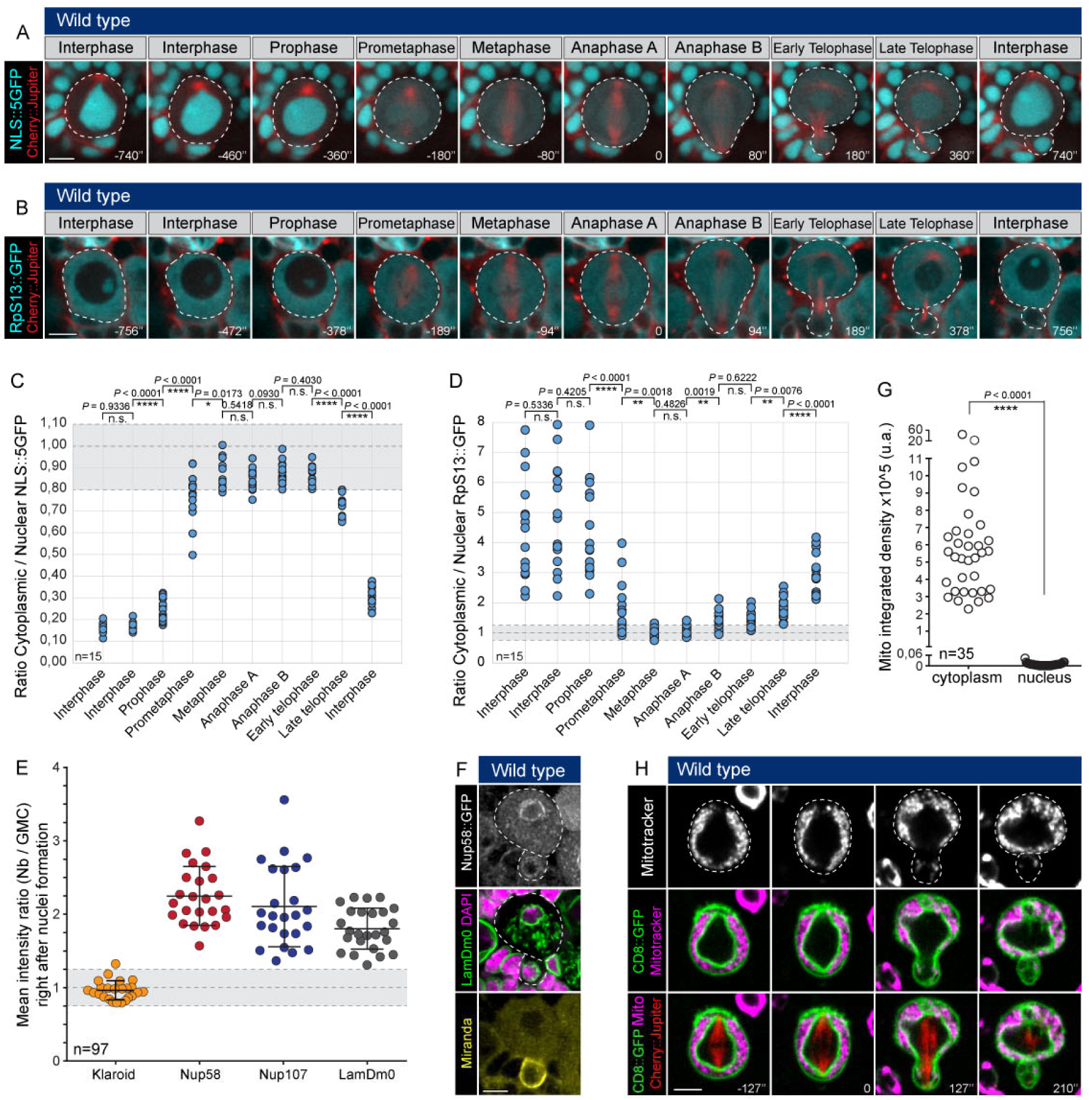
The nuclear envelope of fly neuroblasts is maintained during mitosis and remodelled to generate two sibling nuclei that differ in size and composition. (**A-B**), Time lapse of dividing neuroblast expressing Cherry::Jupiter (red) and (**A**) NLS::5GFP (blue) or (**B**) RpS13::GFP (blue). (**C-D**) Mean intensity of (**C**) NLS::5GFP or (**D**) RpS13::GFP is measured in the cytoplasm and in the nuclear compartment for 10 time points throughout the cell cycle, then the ratio (cytoplasm/nuclear compartment) is plotted on the graph. The grey area corresponds to a ratio equal to 1 ± 0.25. Number of analysed cells = 15 for each condition. Asterisks denote statistical significance, derived from unpaired t tests: n.s.: not significant, *: p ≤ 0.05, **: p ≤ 0.01 and ****: p ≤ 0.0001. (**E**) Graph showing the mean intensity ratio of Klaroid, Nup58, Nup107 and LamDm0 between the neuroblast and GMC nucleus. Bars indicate mean ± standard deviation. Number of analysed cells = 97. (**F**) Representative images of a neuroblast expressing Nup58::GFP (white on the top panel), fixed and stained for LamDm0 (green on the merge), DAPI (pink on the merge) and Miranda (yellow on the bottom panel). (**G**) Graph showing the integrated density of Mitotracker signal in the cytoplasm and in the nuclear compartment, during metaphase. Number of analysed cells = 35. Asterisks denote statistical significance, derived from unpaired t tests: ****: p ≤ 0.0001. (**H**) Time lapse images of dividing neuroblast expressing Cherry::Jupiter (red on the merge), CD8::GFP (green on the merge) and incubated in presence of Mitotracker (white on the top panel, pink on the merges). For each experiment, the data were collected from at least 3 independent experiments. Scale bars are 5µm.

**Figure S2:**
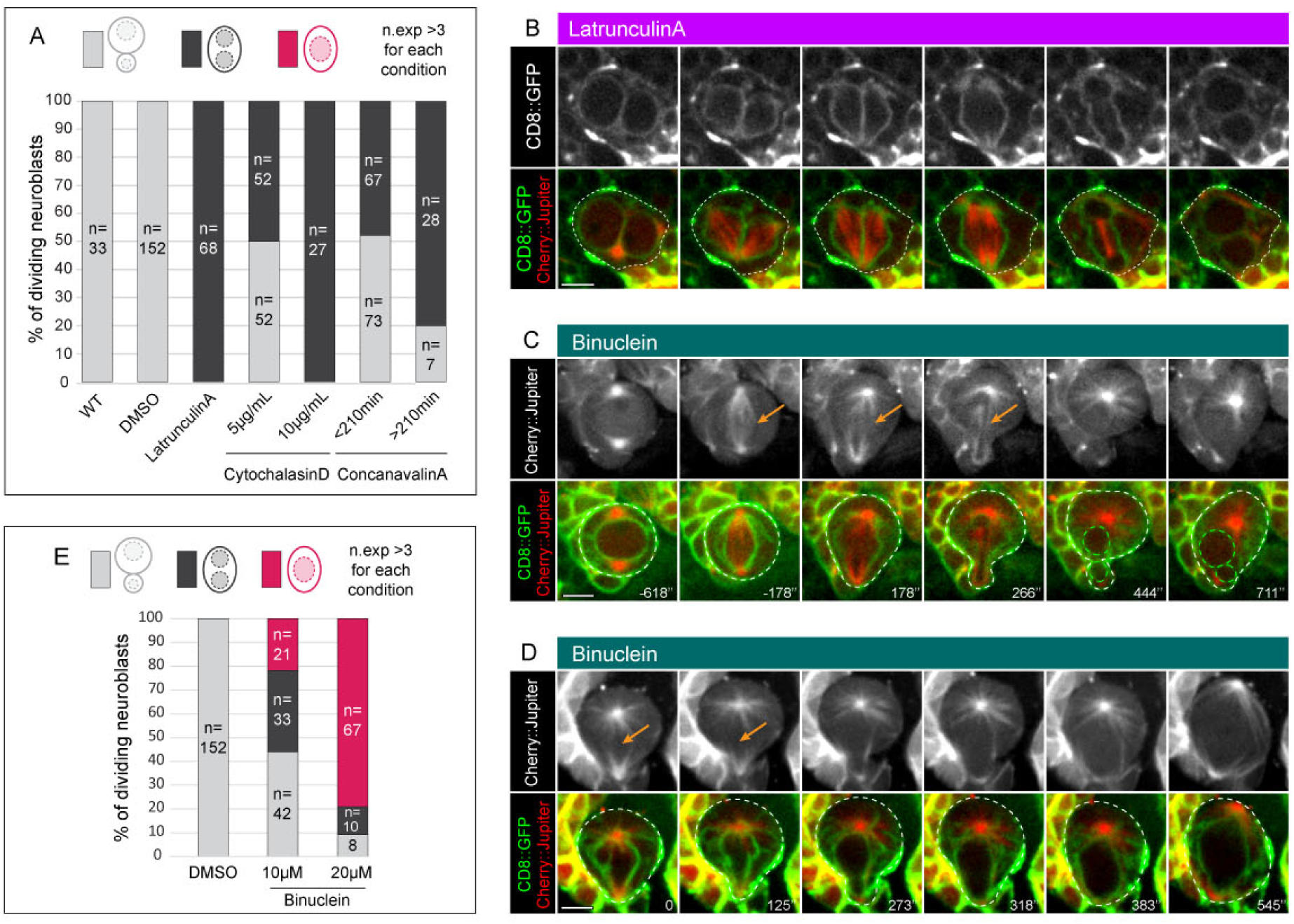
Nuclear division is a sealing-dependent process. (**A**) Quantification of neuroblasts dividing normally (grey category), failing in cell division but still dividing their nucleus (black category) or failing in both cell and nuclear division (pink category). The number of analysed cells for each condition is indicated on the graph. (**B**) Representative timelapse images of binucleated neuroblast expressing CD8::GFP (white on the top panel, green on the merge) and Cherry::Jupiter (red on the merge), failing a second round of cell division. (**C**) Representative timelapse images of Binuclein-treated neuroblast expressing Cherry::Jupiter (white on the top panel, red on the merge) and CD8::GFP (green on the merge) failing only in cell division. Orange arrows show the central spindle. (**D**) Representative timelapse images of Binuclein-treated neuroblast expressing Cherry::Jupiter (white on the top panel, red on the merge) and CD8::GFP (green on the merge) failing in both cell and nuclear division. Orange arrows show the absence of a central spindle. (**E**) Quantification of neuroblasts dividing normally (grey category), failing in cell division but still undergoing nuclear division (black category) or failing in both cell and nuclear division (pink category) after DMSO or Binuclein treatment. The number of cells analysed for each condition is indicated on the graph. For each experiment, the data were collected from at least 3 independent experiments. Scale bar is 5µm.

**Figure S3:**
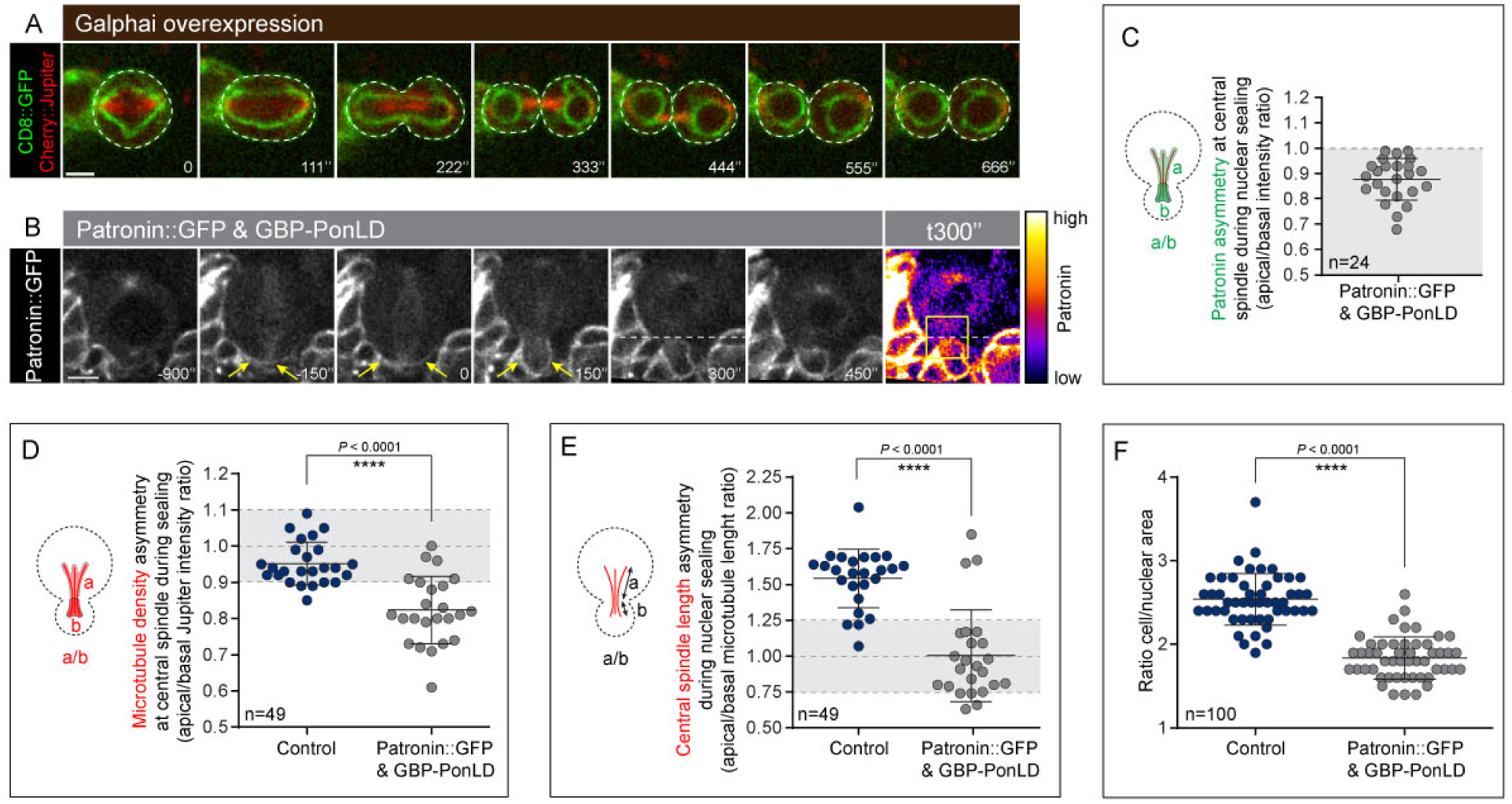
Asymmetric nuclear division depends on nuclear sealing at sites dictated by the spindle. (**A**) Representative time lapse images of a neuroblast overexpressing Galphai, CD8::GFP (green) and Cherry::Jupiter (red). (**B**) Representative time lapse images of a neuroblast expressing Patronin::GFP (white) and the localization domain of Pon fused to GFP Binding Protein (GBP-PonLD). Yellow arrows show the basal recruitment of Patronin::GFP. The last picture corresponds to the sealing time (time point 300”), and is shown with a gradient of colour to highlight the enrichment of Patronin on the basal half of the central spindle. (**C**) Graph showing Patronin distribution at the apical versus basal half of the central spindle. Bars indicate mean ± standard deviation. Number of analysed cells = 24. (**D**) Graph showing the microtubule density ratio of the central spindle (apical versus basal half) in control neuroblasts and in neuroblasts expressing Patronin::GFP and GBP-PonLD. Number of analysed cells = 49. Bars indicate mean ± standard deviation. Asterisks denote statistical significance, derived from unpaired t tests: ****: p ≤ 0.0001. (**E**) Graph showing the microtubule length ratio of the central spindle (apical versus basal half) in control neuroblasts and in neuroblasts expressing Patronin::GFP and GBP-PonLD. Number of analysed cells = 49. Bars indicate mean ± standard deviation. Asterisks denote statistical significance, derived from unpaired t tests: ****: p ≤ 0.0001. (**F**) Graph showing the ratio between cell and nuclear size in control neuroblasts and in neuroblasts expressing Patronin::GFP and GBP-PonLD. Number of analysed cells = 100. Bars indicate mean ± standard deviation. Asterisks denote statistical significance, derived from unpaired t tests: ****: p ≤ 0.0001. For each experiment, the data were collected from at least 3 independent experiments. Scale bars are 5µm.

**Figure S4:**
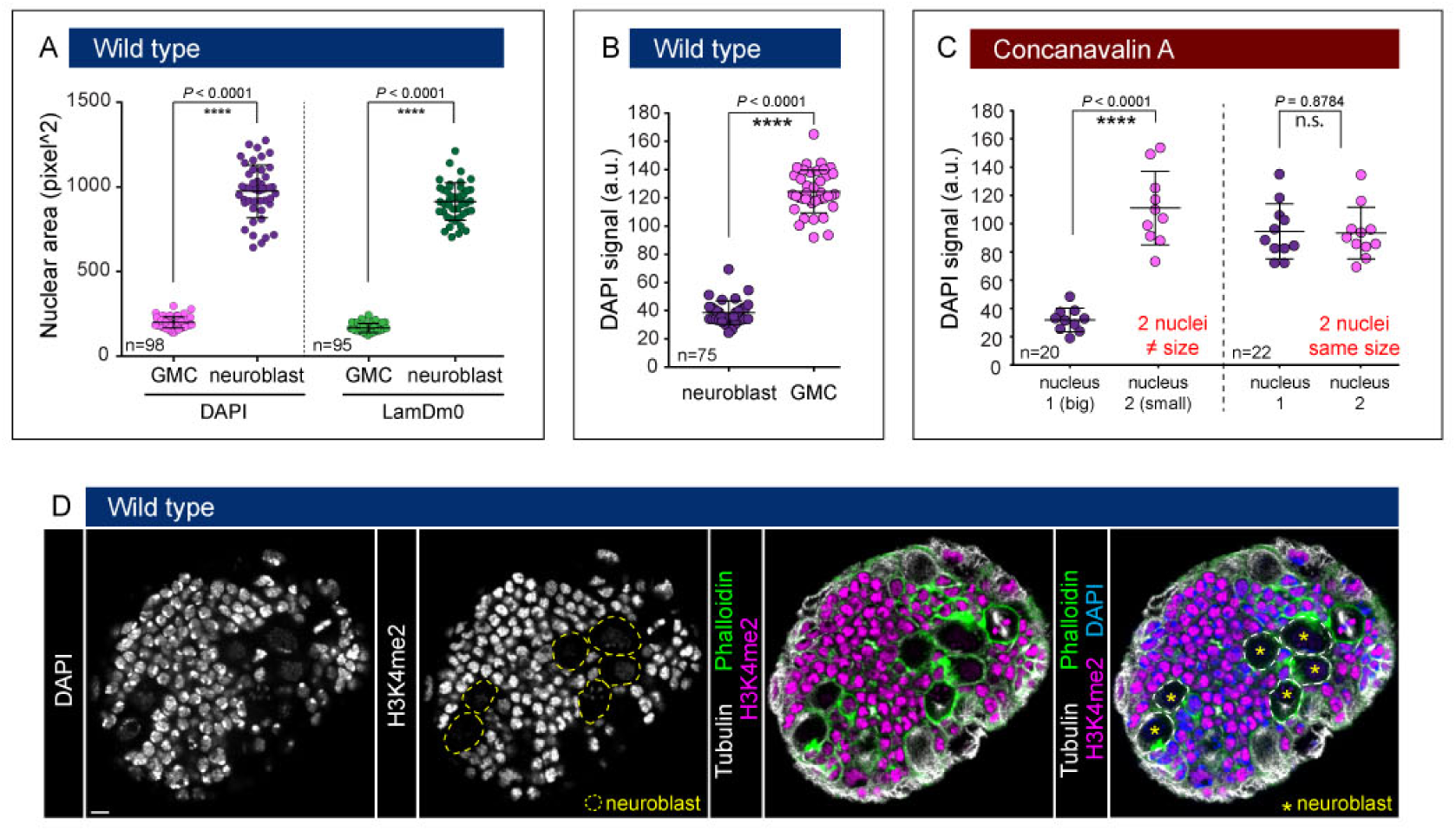
Asymmetries in the nuclear and chromatin organization in neuroblast divisions. (**A**) Graph showing nuclear area of GMC and neuroblast nucleus based on DAPI (left) and LamDm0 (right) staining. Number of analysed cells = 98 and 95 for DAPI and LamDm0 staining, respectively. Bars indicate mean ± standard deviation. Asterisks denote statistical significance, derived from unpaired t tests: ****: p ≤ 0.0001. (**B**) Graph showing the DAPI mean intensity signal in the neuroblast and GMC nucleus. Number of analysed cells = 75. Bars indicate mean ± standard deviation. Asterisks denote statistical significance, derived from unpaired t tests: ****: p ≤ 0.0001. (**C**) Graph showing the DAPI mean intensity signal in sibling nuclei of binucleated neuroblasts. Number of analysed cells = 42. Bars indicate mean ± standard deviation. Asterisks denote statistical significance, derived from unpaired t tests: n.s.: not significant, ****: p ≤ 0.0001. (**D**) Wild type brain lobe fixed and stained for DAPI (white on the left and blue on the merge), H3K4me2 (white on the second panel and pink on the merges), Tubulin (white on the merges) and Phalloidin (green on the merges). Yellow dashed lines and yellow stars represent neuroblasts. For each experiment, the data were collected from at least 3 independent experiments.

**Figure S5:**
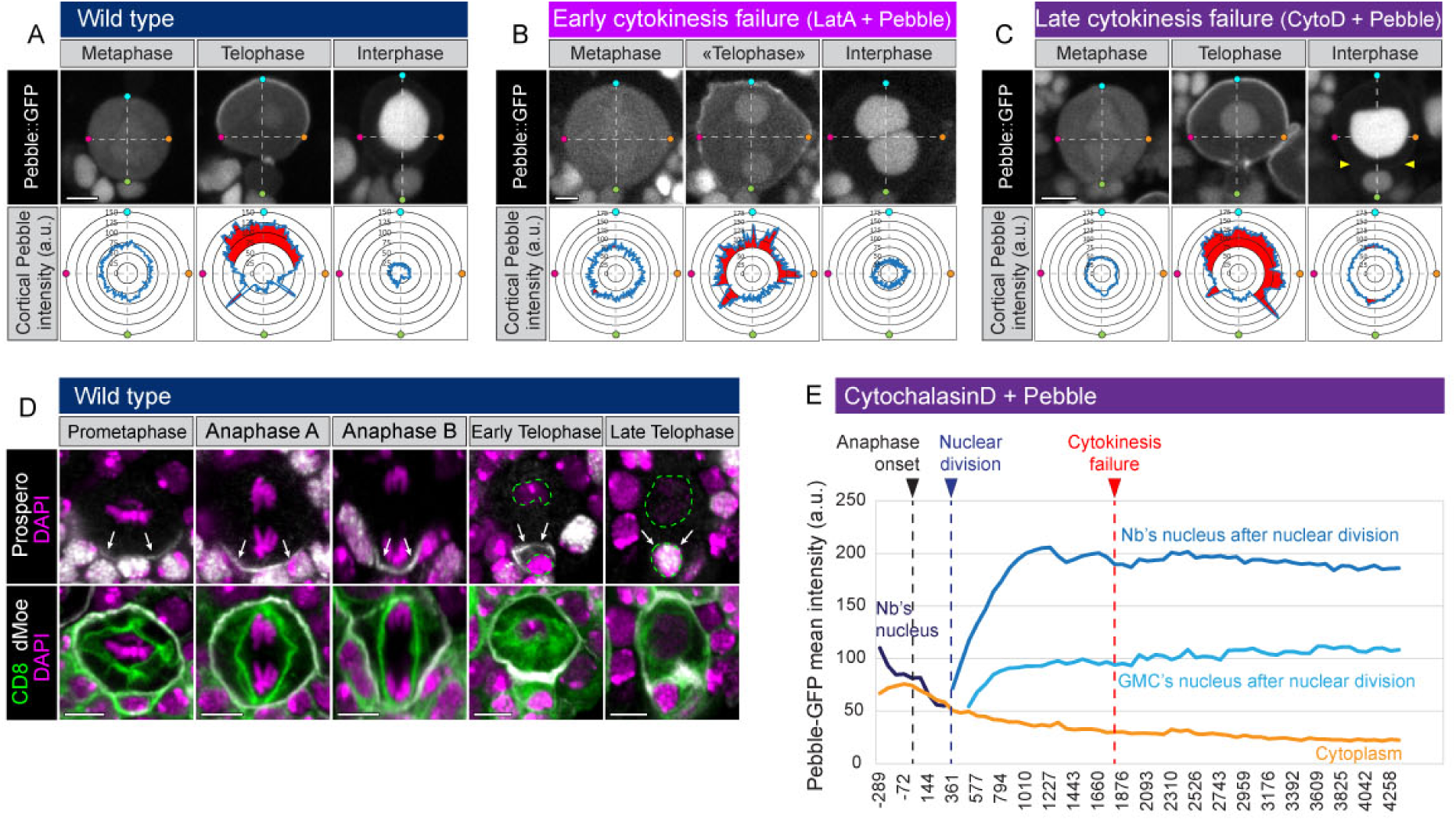
Nuclear identity is set by the relative timing of nuclear division, cell division and cortical release of nuclear proteins. (**A-C**) Pebble::GFP distribution in metaphase, telophase and interphase (white), (**A**) in an untreated neuroblast, (**B**) after early cytokinesis failure (Latrunculin A treatment in combination with Pebble over-expression) and (**C**) after late cytokinesis failure (Cytochalasin D treatment in combination with Pebble over-expression). For each condition and time points (metaphase, telophase and interphase) the Pebble intensity along the cortex is measured and plotted on the rose plots. (**D**), Immunostaining of dividing neuroblasts expressing CD8::GFP (green on the merge) for Prospero (white on the top panel) and for the cortical marker dMoesin (white on the merge). DNA is stained with DAPI (pink). (**E**), Graph showing the Pebble::GFP mean intensity in the cytoplasm (orange curve), as well as along the nuclear membrane before nuclear division (dark blue curve; called “Nb’s nucleus”) and after nuclear division (blue curves corresponding to the neuroblast and GMC nuclei). The vertical black dashed line corresponds to the anaphase onset, the blue one to the nuclear division and the red one to the cytokinesis failure. For each experiment, the data were collected from at least 3 independent experiments. Scale bars are 5µm.

**Video 1**.**avi related to Figure 1**

The nuclear envelope of neuroblasts persists during mitosis and is remodelled to generate sibling nuclei with different size. Neuroblast expressing the plasma and nuclear membranes marker CD8::GFP (white on the left panel, green on the merge) and the spindle marker Cherry::Jupiter (white on the middle panel, red on the merge). Scale bar is 5 µm. Stopwatch shows time in minutes and seconds in regards to anaphase onset.

**Video 2**.**avi related to Figure 1**

The nuclear envelope of neuroblasts retains elements of its identity as neuroblasts pass through mitosis. Neuroblast expressing the Endoplasmic Reticulum luminal reporter protein BIP::SfGFP::HDEL (white on the left panel, green on the merge) and the spindle marker Cherry::Jupiter (white on the middle panel, red on the merge). Scale bar is 5 µm. Stopwatch shows time in minutes and seconds in regards to anaphase onset.

**Video 3**.**avi related to Figure 1**

The nuclear/cytoplasmic diffusion barrier is lost upon entry into mitosis. Neuroblast expressing NLS::5GFP (white on the left panel, blue on the merge) and the spindle marker Cherry::Jupiter (white on the middle panel, red on the merge). NLS::5GFP is leaving the nucleus as neuroblast enters prometaphase. Scale bar is 5 µm. Stopwatch shows time in minutes and seconds in regards to anaphase onset.

**Video 4**.**avi related to Figure 1**

The nuclear/cytoplasmic diffusion barrier is lost upon entry into mitosis. Neuroblast expressing the ribosomal subunit RpS13::GFP (white on the left panel, blue on the merge) and the spindle marker Cherry::Jupiter (white on the middle panel, red on the merge). RpS13::GFP is entering the nucleus as neuroblast enters prometaphase. Scale bar is 5 µm. Stopwatch shows time in minutes and seconds in regards to anaphase onset.

**Video 5**.**avi related to Figure 2**

A lamina persists throughout cell division. Neuroblast expressing LamDm0::GFP (white on the left panel, green on the merge) and Cherry::Jupiter (red on the merge). While the levels of LamDm0::GFP present at the interphase nuclear envelope were visibly reduced during mitosis, some remained at the nuclear envelope throughout. Scale bar is 5 µm. Stopwatch shows time in minutes and seconds in regards to anaphase onset.

**Video 6**.**avi related to Figure 3**

Nuclear division is a sealing-dependent process. Latrunculin-treated neuroblast expressing CD8::GFP (white on the left panel, green on the merge) and the spindle marker Cherry::Jupiter (red on the merge). Nuclear membranes expand to progressively close the nuclear compartment generating daughter nuclei. Scale bar is 5 µm. Stopwatch shows time in minutes and seconds in regards to anaphase onset.

**Video 7**.**avi related to Figure 7**

Control neuroblast expressing Pebble::GFP (white on the left, blue on the merge) and the spindle marker Cherry::Jupiter (red on the merge). Pebble::GFP is asymmetrically recruited to the cortex before to be imported asymmetrically into the sibling nuclei, after cytokinesis. Scale bar is 5 µm. Stopwatch shows time in minutes and seconds in regards to anaphase onset.

**Video 8**.**avi related to Figure 7**

Latrunculin-treated neuroblast expressing Pebble::GFP (white on the left, blue on the merge) and the spindle marker Cherry::Jupiter (red on the merge). Pebble is still asymmetrically recruited at the cortex, before to be imported in a similar way into both daughter nuclei due to an early cytokinesis failure. Scale bar is 5 µm. Stopwatch shows time in minutes and seconds in regards to anaphase onset.

**Video 9**.**avi related to Figure 7**

Cytochalasin-treated neuroblast expressing Pebble::GFP (white on the left, blue on the merge) and the spindle marker Cherry::Jupiter (red on the merge). Pebble is still asymmetrically recruited at the cortex, then asymmetrically imported into sibling nuclei prior late cytokinesis failure. Scale bar is 5 µm. Stopwatch shows time in minutes and seconds in regards to anaphase onset.

